# Silver thiosulfate and Benzyladenine in combination with pruning additively feminizes cassava flowers and modulates transcriptome

**DOI:** 10.1101/2020.12.15.422940

**Authors:** Oluwasanya Deborah, Esan Olayemisi, Hyde Peter, Kulakow Peter, Setter Tim

## Abstract

Cassava, a tropical storage-root crop, is a major source of food security for millions in the tropics. Cassava breeding however is hindered by the poor development of flowers and female flowers in particular, since flower development is strongly skewed towards male flowers. Our objectives were to test plant growth regulator and pruning treatments for their effectiveness in field conditions in improving flower production and fruit set in cassava. Pruning the fork type branches that arise at the shoot apex immediately below newly formed inflorescences stimulated inflorescence and floral development. The anti-ethylene silver thiosulfate (STS) also increased flower abundance. Both pruning and STS increased flower numbers without influencing sex ratios. In contrast, the cytokinin benzyladenine (BA) feminized flowers without increasing flower abundance. Combining pruning and STS treatments led to an additive increase in flower abundance; with the addition of BA, over 80% of flowers were females. This three-way treatment combination of pruning+STS+BA also led to an increase in fruit development. Transcriptomic analysis of gene expression in tissues of the apical region and developing inflorescence revealed that the enhancement of female flower development by STS+BA was accompanied by the downregulation in of several genes associated with repression of flowering, including Tempranillo 1 (TEM1), GA receptor GID1b, and ABA signaling genes ABI1 and PP2CA. We conclude that treatments with pruning, STS and BA create widespread changes on the network of hormone signaling and regulatory factors beyond ethylene and cytokinin.

## 1 Introduction

Cassava (*Manihot esculenta*) is a perennial tropical plant of the Euphorbiaceae family cultivated as an annual crop for its starchy storage roots (Liu et al. 2014). It constitutes an important source of calories for over 750 million people (Tuteja et al. 2012). It is also a source of starch with expanding potential for industrial applications (Li et al. 2017). Continual crop improvement is required to ensure efficient cassava production to meet growing needs from an increasing population matched with changing environments globally. Cassava improvement has recently received renewed attention with major projects to investigate the potential use of genomic selection in breeding (Wolfe et al. 2017), detailed studies into its source-sink relationships (Sonnewald et al. 2020) and investigations to improve its photosynthetic efficiency (De Souza and Long 2018). These improvement efforts are geared towards small holder farmers with interest in translating cassava end product quality traits into breeding outcomes (Iragaba 2019). Although cassava can be clonally propagated by stem cuttings, achieving crop improvement requires genetic hybridization and associated recombination, and hence the need for a means of sexual propagation that is timely and synchronous amongst a diverse genetic population.

In the Euphorbiaceae family, sexual reproductive structures are referred to as cyathia, which is a modification that has provided this family the advantage of shifting from wind pollination to insect pollination (Horn et al. 2012). A single cassava male cyathium is comprised of multiple reduced stamens (Perera et al. 2013) while a female cyathium possesses a trilocular ovary, meaning that upon pollination, fruits are capable of producing three seeds (Nassar 1980). Female and male cyathia are borne on the same inflorescence separately. For ease of description, cyathia in this study will be referred to as flowers. Inflorescences and associated flowers are developed from the shoot apical meristem. Following floral initiation, the two to three buds beneath the inflorescence develop into branches forming a fork. Fork type branches each bear new shoot apical meristems in sympodial growth which after some growth form branches and inflorescences, in turn, at tier 2, tier 3, etc. Perera et al. (2013) provides a botanical description of cassava reproductive structures.

There are five bottlenecks with cassava’s reproductive development that challenge breeding. These are (1) late flowering (or in some cases no flowering at all) of certain genotypes with traits of interest, (2) premature abortion of flowers before anthesis, (3) disproportionate number of male flowers and in some cases no female flowers, in cases where flowers are not aborted, (4) non synchronous development of flowers, (5) low probability of fruit development even when flowering and pollination are successful (Adeyemo et al. 2017; Hyde et al. 2020; Ceballos et al. 2004; Ceballos et al. 2016; Halsey et al. 2008; Souza et al. 2020).

Investigations into the reproductive biology of cassava has led to the development of several potential interventions to improve reproductive performance in cassava: transgenic intervention by overexpressing Flowering Locus FT (Adeyemo et al. 2017; Odipio et al. 2020; Bull et al. 2017), modulating photoperiod and temperature to accelerate flowering time (Adeyemo et al. 2019), application of silver thiosulfate to enhance flower development and longevity (STS) (Hyde et al. 2020), or pruning young subtending branches to alleviate flower abortion (Pineda et al. 2020). None so far has focused on increasing the proportion of female flowers. Female flowers are critical to cassava breeding; however, flowers on cassava inflorescences are typically about 90% male, and each female flower produces only three seeds, at most. In other members of the Euphorbiaceae family, synthetic cytokinin, Benzyladenine, has been used to increase female flower numbers and fruits (Fröschle et al. 2017; Fu et al. 2014; Pan and Xu 2011).

To increase the proportion of female flowers in cassava we tested the effect of STS, BA and pruning, singly and combined, on the first flowering event (i.e., flowering at tier 1) under field conditions in Nigeria. We also analyzed the transcriptome of young apical tissues in control plants and in plants receiving either pruning or STS + BA, or the combined set of treatments. We specifically studied the expression pattern of differentially expressed genes relevant to hormone signaling and flower development. Our findings indicate that STS or pruning increase flower numbers over the control but have no effect on the female to male ratios. While BA is able to completely feminize flowers, it does not affect the number of total flowers. Combining BA with either STS or pruning or both allows for development of predominantly female flowers on a larger population of flowers. The transcriptome changes under STS+BA treatment indicates effects on other hormone signaling beyond ethylene and cytokinin.

## 2 Materials and methods

### 2.1 Plant materials

Three genotypes, representing three flowering times and extents of flower prolificacy were used for plant growth regulator (PGR) studies. These were IITA-TMS-IBA980002 (early and profuse), IITA-TMS-IBA30572 (middle genotype) and TMEB419 (late and poor flowering). In this paper these genotypes will be referred to as 0002, 30572, and 419, respectively. 0002 was used to optimize the method of plant growth regulator (PGR) application in 2017. This experiment was conducted between June and December of 2017. Experiments examining the effect of PGRs on female flower development were conducted between June and December of 2018 and 2019.

### 2.2 Field conditions

All experiments for phenotyping were conducted under field conditions at the International Institute of Tropical Agriculture (IITA), Oyo State, Ibadan (7.4° N and 3.9°E, 230m asl). The soil was an Alfisol (oxicpaleustalf) (Moormann et al. 1975). The land was tilled and ridged with 1 m spacing; plants were sown on top of the ridge. The land in 2017 in Experiment I was previously planted with yam while the field in 2018 and 2019 in Experiment II was previously planted with maize; no extra nutrients or soil amendments were added to the soil. Fields were kept free of weeds with hand weeding.

### 2.3 Plant growth regulators and method of application

Silver thiosulfate (STS) was prepared by mixing 1 part 0.1 M silver nitrate (AgNO_3_) dropwise with 4 parts 0.1M sodium thiosulfate (Na_2_S_2_O_3_), yielding a 20 mM stock solution. The stock solution was diluted with distilled water to the required concentration as specified in each experiment.

Benzyladenine (BA) solution was prepared by diluting a 1.9% (w/v) BA stock (MaxCel®, Valent BioSciences Corporation, Libertyville, IL, USA) with distilled water to respective concentrations. PGRs were applied either by spray (about 5 mL) to the shoot apex, every seven days or by the “petiole feeding” method every 14 days. In the petiole feeding method, the leaf blade was removed using a surgical scissors and the petiole was inserted into a 15-mL conical-bottom centrifuge tube (Falcon Brand, Corning, NY, USA) containing 5 – 10 mL of PGR solution. PGR was taken up via the petiole into xylem from which it was distributed internally to target organs in the leaves and apex (Figure 1a). Petioles were allowed to remain immersed in the PGR solution for 72h after which tubes were removed. On weeks with petiole feeding, spray treatments were applied 24 h after petiole treatments. PGR treatments were initiated six weeks after planting.

**Figure 1.**
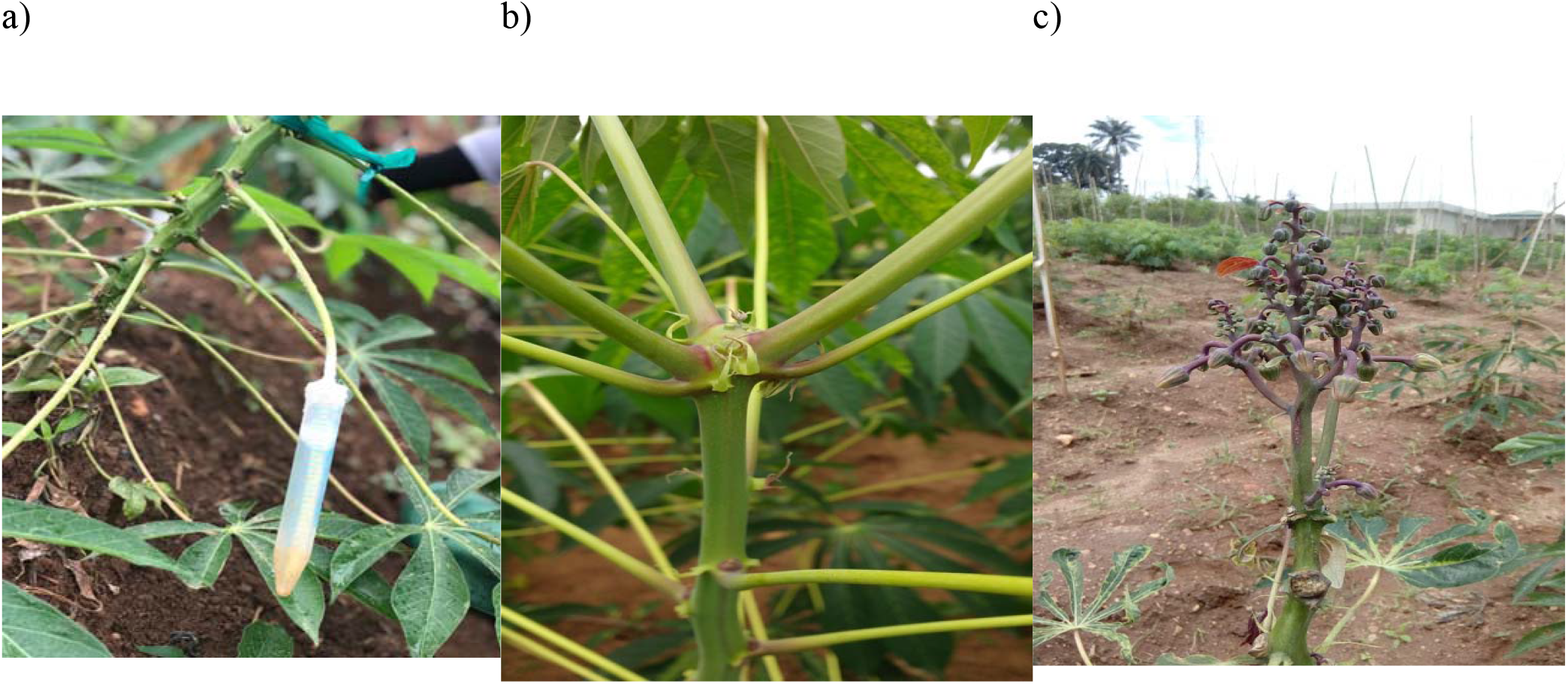
Photos of cassava a) receiving treatment by petiole feeding b) with unpruned fork-type branches; inflorescence aborting c) pruned fork-type branches with thriving flowers

### 2.4 Pruning reproductive branches

Shoot apexes were inspected weekly using headband magnifier glasses (10X magnification) to identify and score forking events. In plants that had forked, young reproductive branches, 2 cm or smaller, were excised carefully without damage to inflorescence using surgical blades, as previously described by Pineda et al. (2020). Lateral branches which developed about 10 cm below the shoot apex were excised periodically until fruit development. Photos of pruned and unpruned plants are shown in (Figure 1 b,c)

### 2.5 Flower data collection

The number of male and female flowers and fruits were counted for each plant weekly and recorded using Field Book computer application (Rife and Poland 2014). Data was collected for at least 20 weeks. For analysis, the maximum flower or fruit count for each plant over the 20-week period was used to represent response to treatment.

### 2.6 PGR Experiment I: Application method optimization

A matrix of 100 plants of IITA-TMS-IBA980002 were grown in a 10 x 10 arrangement at a planting distance of 1 m ×1 m. The matrix was divided into 10 plots to allow for treatment assignments. At six weeks after planting, 10 PGR treatments were randomly assigned to each plot, with each plant as the experimental unit. Treatments were various combinations of petiole-fed STS or H_2_O with spray-applied BA at four flower developmental stages. To determine the effect due to STS alone, STS or H_2_O were applied by petiole feeding in combination with H_2_O sprayed to the shoot apex. Five mL of 8 mM STS was used for the first treatment then reduced to 5 mL of 4 mM STS for subsequent treatments to limit phytotoxicity. Concentration of BA used was 0.22 mM. Treatments are summarized in Table 1.

**Table 1.** BA application treatment summary for Experiment I.

| Apical spray | Flower Developmental Stage |
| --- | --- |
| Control | Spray water to shoot apex throughout time frame |
| BA_Always | Spray BA beginning before flower appearance and continue until end of experiment |
| BA_Early | Spray BA beginning before flower appearance and stop at flower appearance |
| BA_Mid | Spray BA beginning at flower appearance and stop when first female flower antheses |
| BA_Late | Spray BA beginning at 21 d after flower appearance and continue until end of experiment |

### 2.7 PGR Experiment II: PGR and pruning effect on female flower development

In 2018 and 2019, experiments were conducted using a split-split-split plot design. Each experiment comprised six plots, each of which were split into three subplots each with one of the genotypes. Each genotype subplot was split into 5 PGR treatments (as shown in Table 2) and finally each treatment was split into two pruning levels – pruned versus unpruned. In 2018, a total of 1440 plants were sown, while in 2019, 720 plants were sown. Petiole feeding of STS and BA was with either 10 mL of 2 mM STS (5 mL of 4 mM STS was used in the first month of treatment in 2018), 5 mL of 0.5 mM BA, or a mixture of 5 mL of 2 mM STS and 0.125 mM BA. Spray treatments of BA were applied to the shoot apex with 0.5 mM BA.

**Table 2.** Summary of PGR treatments for Experiment II.

| PGR | BA | BA | STS |
| --- | --- | --- | --- |
| Method of delivery: | apical spray | petiole fed | petiole fed |
| **Treatment name:** |  |  |  |
| **Control** |  |  |  |
| **BA** | x |  |  |
| **BA+BA** | x | x |  |
| **STS+BA** | x |  | x |
| **STS+BA+BA** | x | x | x |

### 2.8 Statistical Analysis

Count data, such as the numbers of total flowers, female flowers, and fruits were modelled using the negative binomial model while ratios such as the proportion of total flowers that were female and proportion of female flowers that set fruit were modelled using the binomial model. Due to similarities of genotypic response to PGR and pruning (different magnitudes of changes but similar trends), the means of genotypes used in 2018 and 2019 are reported here.

Models were built using the package called Generalized Linear Mixed Models using Template Model Builder (glmmTMB) (Brooks et al. 2017) in R (Team 2013). Sources of variation for the experiment were STS treatment (+STS, -STS) and BA timing, and the interaction between STS treatment and BA timing. Plots were random effects. For Experiment II, sources of variation were PGR treatment, pruning and the interaction between PGR treatment and pruning. Year was analyzed as a fixed effect while plot as a random effect. The emmeans package (Lenth 2019) was used for post-hoc tests. Multiple means comparison was carried out using the Tukey-HSD method.

### 2.9 Transcriptomic Analyses

IITA-TMS-IBA980002 was grown in a greenhouse at the Guterman Bioclimatic laboratory (Cornell University, Ithaca, NY, USA) as described by Hyde et al (2020) and exposed to one of four treatment combinations (a 2 × 2 matrix of treatments): a) either control or pruned at first inflorescence appearance, and b) either a control or PGRs with a combination of petiole-fed STS, and apex-sprayed BA. These were applied to plants at tier 1 of fork branching. Young inflorescence tissue, about 0.25 – 0.5 cm, comprising the shoot apex and some bracts but excluding fork-type branches were harvested from control, pruned and PGR treated plants. Samples were collected four days after pruning or at a similar developmental stage in unpruned plants.

Total RNA was extracted from each sample by a modified CTAB protocol. Samples were ground to a fine powder in a mortar and pestle chilled with liquid N_2_; about 0.15 mL of the powder was vigorously mixed for 5 min with 0.4 mL of CTAB extraction buffer (1% [w/v] CTAB detergent, 100 mM Tris-HCl [pH 8.0], 1.4 M NaCl, 20 mM EDTA, and 2% [v/v] 2-mercaptoethanol); 0.2 mL of chloroform was added and mixed for 1 min, tubes were centrifuged at 14,000 g for 10 min and 200 μL of the top layer was removed to a new tube. To these samples was added 700 μL of Guanidine buffer (4M guanidine thiocyanate, 10 mM MOPS, pH 6.7) and 500 μL of ethanol (100%). This mixture was applied to silica RNA columns (RNA mini spin column, Epoch Life Science, Missouri City, TX, USA), then washed with 750 μL of 1) MOPS-ethanol buffer (10 mM MOPS-HCl [pH 6.7], 1 mM EDTA, containing 80% [v/v] ethanol), 2) 80% ethanol (twice), and 3) 10 μL RNAase-free water to elute the RNA (twice). The RNA quality was evaluated with a gel system (TapeStation 2200, Agilent Technologies, Santa Clara, CA, USA).

Differential Gene expression analysis was conducted using the DESeq2 package by Bioconductor (Love et al. 2014). Each transcript was annotated by the best match between Manihot esculenta genome v7 and the Arabidopsis genome as presented at Phytozome13 (Goodstein et al. 2012). Gene ontology and enrichment analysis were carried out using the ShinyGO app (http://bioinformatics.sdstate.edu/go/) (Ge et al. 2020).

A combined list of Arabidopsis flowering genes were obtained from the Max Planck Institute (https://www.mpipz.mpg.de/14637/Arabidopsis_flowering_genes) and Flowering Interactive Database (FLOR-ID) (http://www.phytosystems.ulg.ac.be/florid/) (Bouché et al. 2016); a list of hormone signaling genes sourced through the Database for Annotation, Visualization and Integrated Discovery (DAVID) (https://david.ncifcrf.gov/) (Dennis et al. 2003) and a list of Cassava MADS-Box MIKC genes were obtained from iTAK data-base (http://itak.feilab.net/cgi-bin/itak/db_family_gene_list.cgi?acc=MADS-MIKC&plant=3983) (Zheng et al. 2016) were used to determine expression pattern of genes by flowering, hormone signaling and MADS-Box categories.

## 3 Results

### 3.1 PGR Experiment I: Application method optimization

In previous studies under greenhouse growth conditions, STS spray treatments increased flower numbers and longevity (Hyde et al. 2020); however, when sprayed onto leaves, cassava foliage sometimes suffered phytotoxicity. It was possible to substantially decrease the quantity of STS used with similar benefit if the spray was localized to the young folded leaves and region of the shoot apical meristem. However, when these PGRs have been used in preliminary studies on cassava with spray application in the field, their effectiveness has not been clear cut (Setter, personal communication). A new petiole feeding method was developed, similar to that reported by Lin et al. (2011). This method introduced more modest quantities of PGR internally via xylem of the petiole and delivered these substances to the shoot such that phytotoxicity was decreased (Hyde and Setter personal communication). In the current studies we used petiole feeding as a method of STS application in field studies. Further, we investigated whether BA applied as a spray to the immature tissues, affects flower development as a sole treatment or when combined with STS delivered through the petiole.

Compared to the control without any PGR treatment, STS significantly (P≤0.05) increased total number of flowers as a sole treatment, and in combination with BA (Figure 2a). STS treatment increased both the number of female flowers (Figure 2b) and male flowers. BA treatments did not affect the total number of flowers (Figure 2a); however, in comparison with the control not treated with STS, when BA was applied as a sole treatment throughout flower development (BA_Always) or at the early stage (BA_Early), it created a weakly significant (P≤0.10) increase in the number of female flowers (Figure 2b). These increases were associated with a unique effect of BA on increasing the proportion of total flowers that were female (Figure 2d). In this case the interaction between STS and BA treatments was significant. Early and continuous BA treatments (with or without STS) significantly increased the fraction for female flowers relative to the controls. Later BA treatments (Mid or Late) did not significantly affect the fraction of female flowers. STS alone had a slightly higher fraction of flowers that were female (Figure 2d), but it significantly (P≤0.05) increased fruit numbers (Figure 2c). In contrast, BA did not affect fruit numbers. For total flower, female flower and fruit numbers the interaction between STS and BA treatment at different developmental stages was not significant (P<0.05).

**Figure 2.**
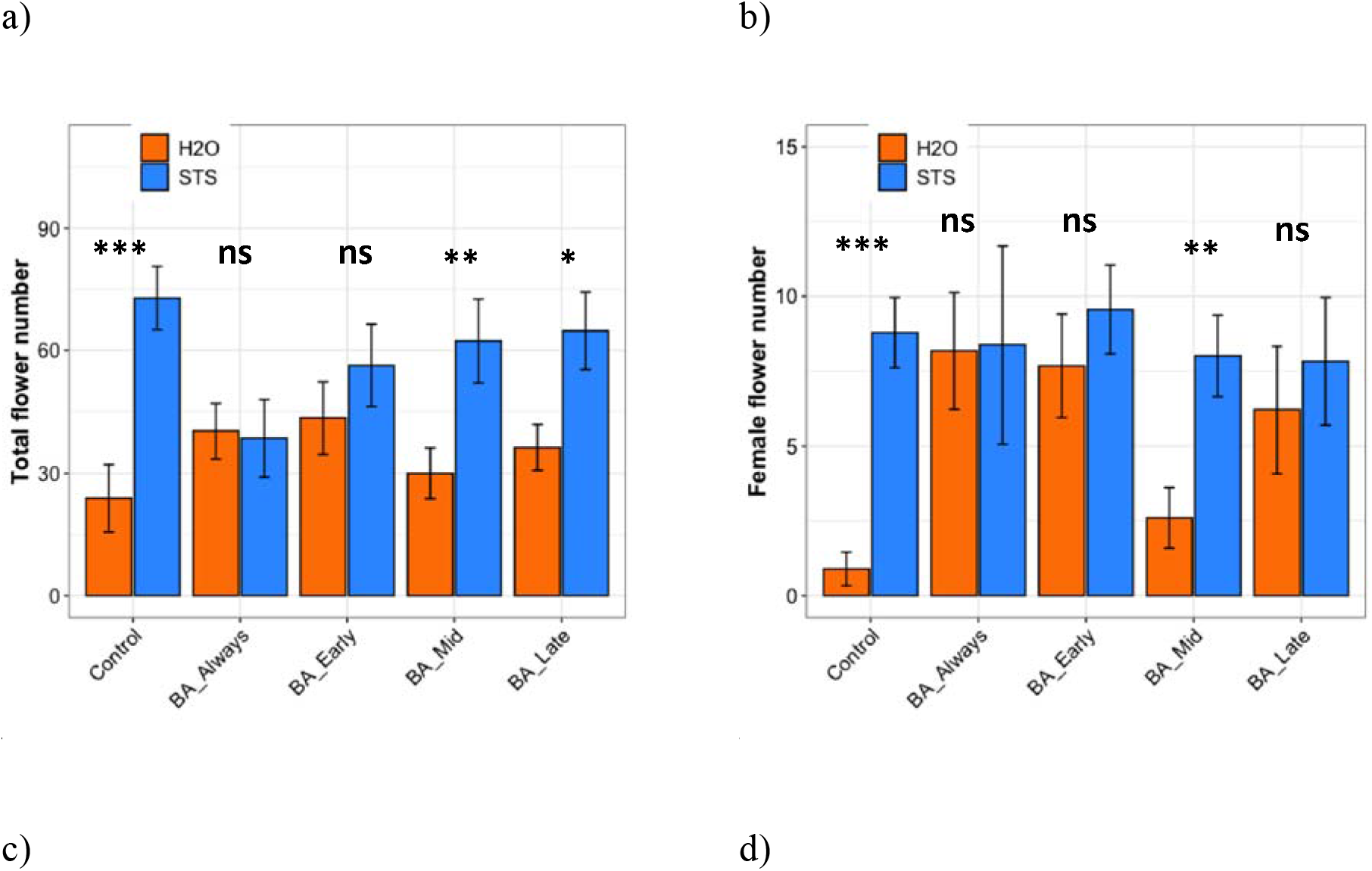
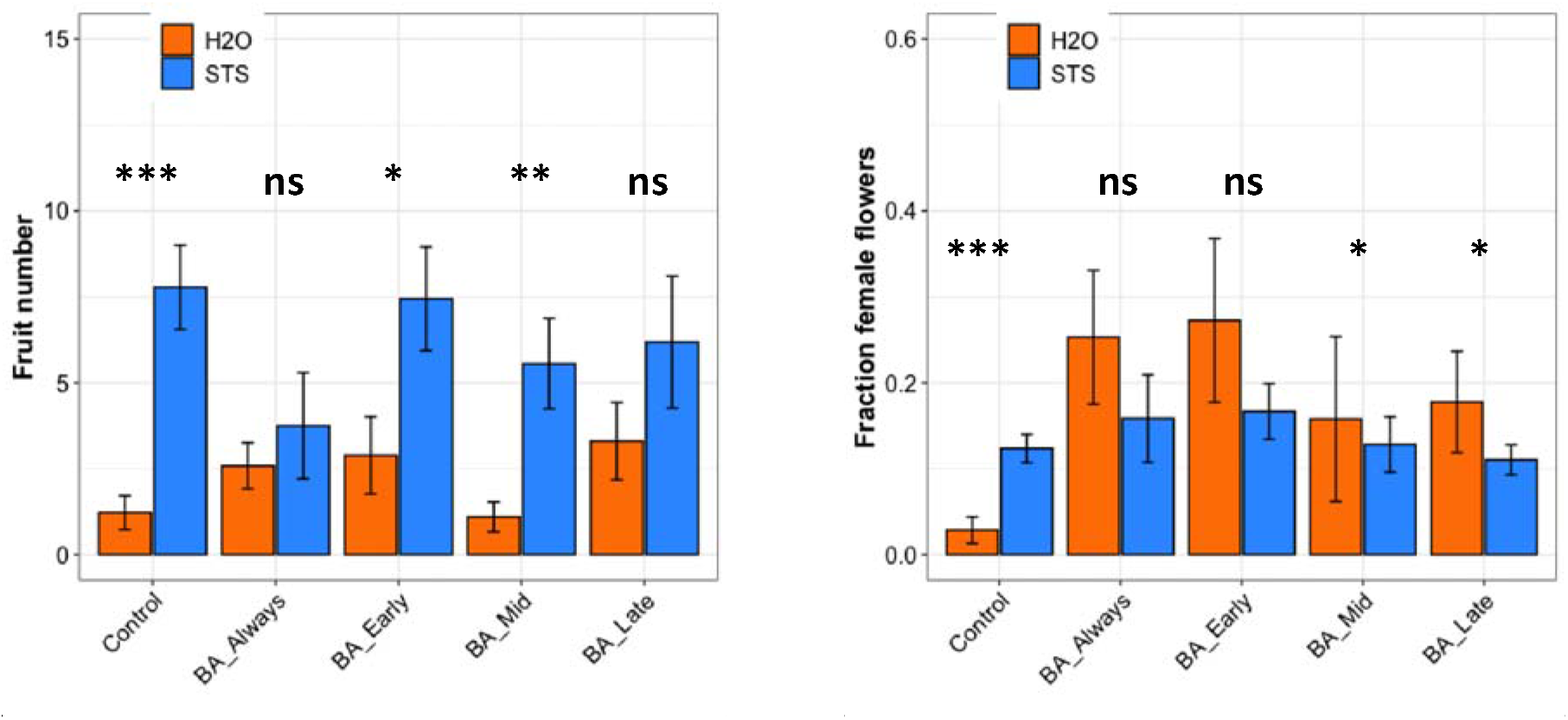
Effect of STS and BA treatments on: a) total flower numbers per plant, b) number of female flowers per plant, c) number of fruit per plant, and d) fraction of female flowers. STS versus no-STS (H_2_O) treatment is indicated by bar color as shown in the legends. BA treatments were applied thoughout floral development (BA_Always), or they were initiated at three timings (BA_Early, BA_Mid, or BA_Late). The study was conducted using genotype IITA-TMS-I980002 in the field at Ibadan, Nigeria in 2017. See M&M Section for details. Data shown is the mean ± SE of 10 replicates; asterisk indicate statistical significance in a pairwise comparison between +STS and no STS; P<0.05 (*), P<0.01 (**), P<0.001 (***).

### 3.2 PGR Experiment II: PGR and pruning effect on female flower development

From Experiment I, above, we obtained evidence that STS treatment increases flower and fruit numbers and that spraying BA to the shoot apex feminized flowers and increased the proportion of flowers that were female. In Experiment II, we tested two additional factors. First is the potential effect on flowering of pruning branch shoots that arise just below the terminal meristem where inflorescences initiated (Pineda et al. 2020). Second is the potential benefit of including BA in petiole feeding rather than only via external spray to leaves and/or apical regions. We therefore investigated the interaction between continuous BA sprays with 1) petiole-fed BA, 2) petiole-fed STS and 3) a mixture of STS and BA (see Materials and Methods). We also increased the BA concentration to 0.5mM and studied PGR effects with or without pruning (see Materials and Methods). The effect on individual genotypes are shown in supplementary figure 1, while averaged, are presented below.

#### Total flower number

In the absence of pruning and STS, total flower numbers in BA-only treatments – whether applied by spray (BA) or via the combination of apical spray and petiole methods (BA+BA) – were not significantly different from the control (Figure 3a). In contrast, STS inclusive treatments (i.e. STS+BA and STS+BA+BA) in the absence of pruning significantly increased female flower numbers relative to the control. In the presence of pruning, the total number of flowers increased significantly in all PGR treatments and in the no-PGR pruning treatment relative to their unpruned counterparts. Pruning and STS had an additive effect, as this combination had the largest number of flowers (Figure 3a).

**Figure 3.**
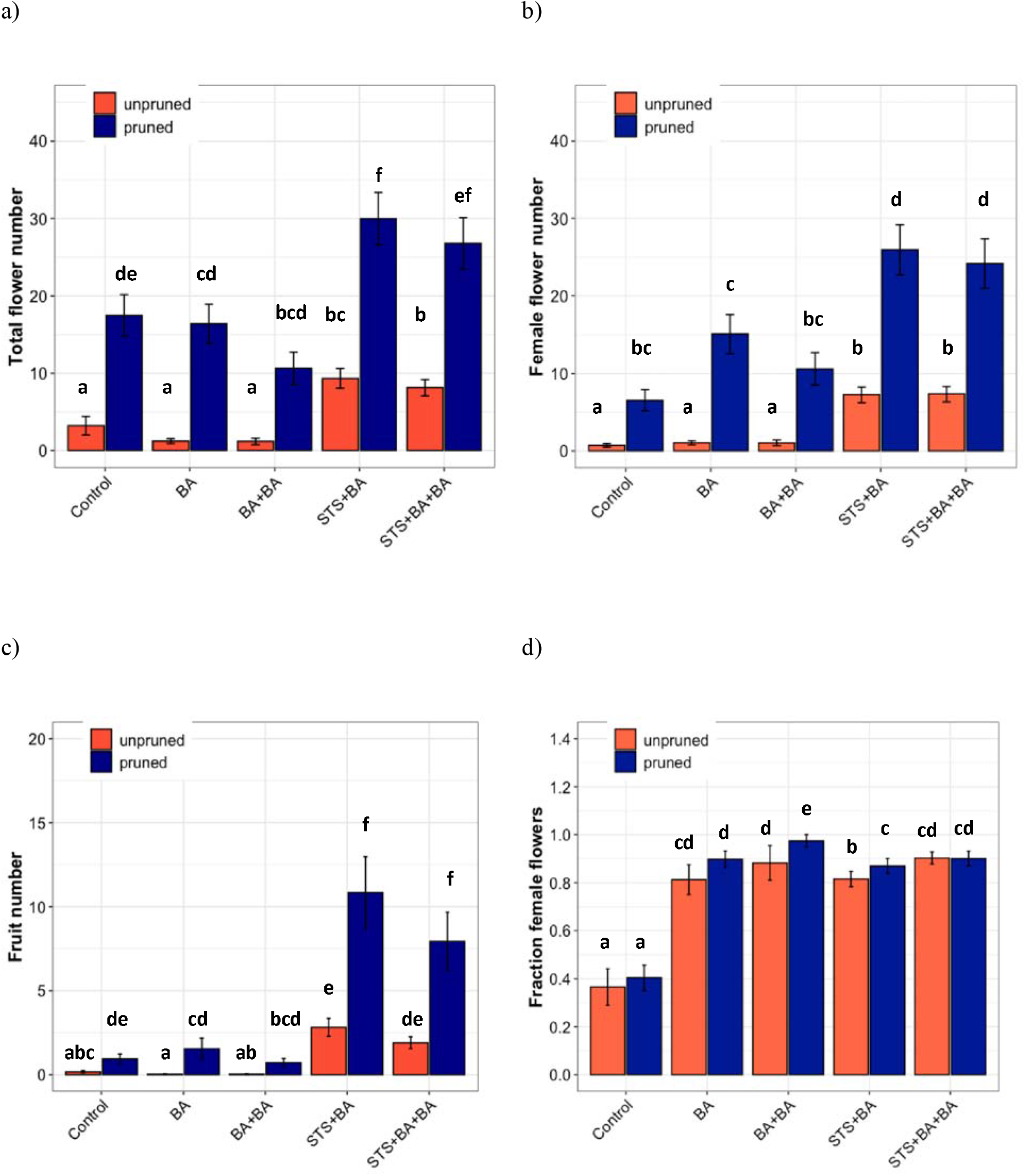
The effect of PGRs and pruning on flower development. a) Total flowers, b) female flower numbers c) fruit numbers d) proportion of female flowers (reflecting only plants with at least one flower). Data shown is the mean of studies conducted in two years (2018 and 2019), each with 6 blocks containing all treatments and 8 (2018) or 4 (2019) plants as experimental units. Treatments with different lowercase letters are significantly (P<0.05) different using Tukey’s HSD test.

#### Female flower number

The effect of treatments on female flower numbers was similar to effects on total flower numbers. BA-only PGR treatments (BA, BA+BA) were not significantly different from the no-PGR treatments without pruning and with pruning (Figure 3b). STS-inclusive treatments (STS+BA, STS+BA+BA) without pruning had significantly more female flowers than BA-only treatments and the no-PGR control without pruning but was equivalent to BA+BA and control with pruning. STS-inclusive treatments with pruning had the highest number of female flowers. (Figure 3b).

#### Fruit numbers

As with other traits, fruit numbers in BA-only treatments were not significantly different from control in the absence or presence of pruning (Figure 3c). STS-inclusive treatments, however, increased fruit numbers relative to the control in both the unpruned and pruned plants such that the combination of pruning and STS yielded largest number of fruits.

#### Female proportion of total flowers

In the controls, female flowers represented 35 to 40% of the flowers in both un-pruned and pruned plants, with the remainder male flowers (Figure 3d). In contrast, all treatments that included BA were significantly different from the controls and had over 80% females, with and without pruning, and with and without STS. Only plants that had flowers were included in this analysis (plants with no flowers were excluded).

### 3.3 Transcriptomics

To advance our understanding of PGR and pruning effects on flowering regulatory processes, we analyzed gene expression in response to PGR and pruning treatments. For this work, we used treatments which had the largest effect in the field: a) STS+BA without pruning, b) STS+BA with pruning, c) pruning without PGR treatment, and d) control (no PGRs and no pruning). This study was conducted on the model genotype 0002, in a controlled environment green house.

#### 3.3.1 Green house phenotype validation

To validate treatment responses that were observed in the field with those in a controlled environment greenhouse, we evaluated flowering traits with select treatments in the greenhouse. Similar to the field study, the controls initiated inflorescences, but did not produce any mature flowers, while pruning or STS+BA as sole treatments produced a modest, number of flowers (Figure 4a); in the pruning treatment all the flowers were male, but in STS+BA, about 80% were female (Figure 4 b,c). Combining STS+BA with pruning increased total and female flower numbers by about three-fold compared to the PGR-or pruning-only treatments. As with field studies, BA-containing treatments increased the number of female flowers and the proportion of flowers that were female (Figure 4 b,c). These findings confirmed a consistent response to treatments under field and greenhouse conditions.

**Figure 4.**
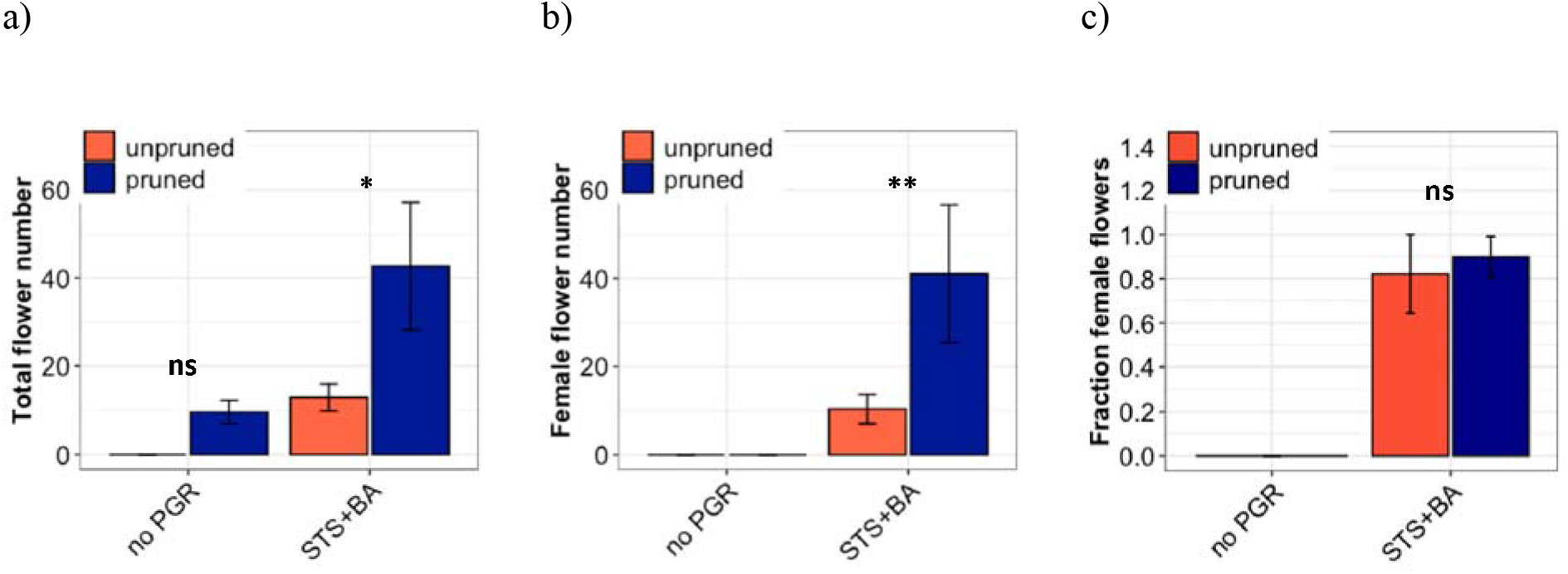
The effect of PGRs (combined STS+BA) and pruning on flower development under greenhouse conditions. a) total flowers, b) female flowers, and c) proportion of female flowers. Data shown is the mean 3 plants per treatment per pruning level for the genotype 0002. * indicates statistical significance on a pairwise comparison of pruned versus unpruned (P<0.05).

#### 3.3.2 Identification of differentially expressed genes and enrichment analysis

Transcriptome analysis was conducted for tissues of the shoot apical region and proceeded in three phases (i) examining transcriptome changes due to pruning alone (design = ~pruning), (ii) examining transcriptome changes due to PGR alone (design = ~pgr), (iii) examining transcriptome changes due to the combination of pruning and PGR (design = ~pruning + pgr). This analysis revealed that the majority of the changes in the transcriptome was due the influence of the PGR treatments. Principle component (PC) analysis indicated that expression was not clearly grouped according to pruned versus unpruned treatments but was clearly grouped according to PGR-treated versus PGR untreated samples (Figure 5a). This grouping was along the first PC axis, which accounted for 54% of the variance. It appeared from the PC analysis that pruning had an intermediate effect in the positive direction along the first principal component whereas PGR had a more substantial effect in the same direction along PC1 axis; the combination of PGR and pruning gave the largest effect in the positive direction of PC1 axis. Analysis of the full model including pruning and PGR revealed that 5440 genes were differentially expressed at a 5% false discovery rate (FDR) correction; 2448 genes were up regulated while 2952 genes were down regulated. Functional analysis revealed that in the PGR-treated versus controls, PGR-upregulated genes were enriched in pathways involving cell proliferation, cell maintenance, and biosynthetic processes; while PGR-down regulated genes were enriched in pathways involved in plant hormone signal transduction, photosynthesis and degradation metabolism, among others (Figure 6 a,b).

**Figure 5.**
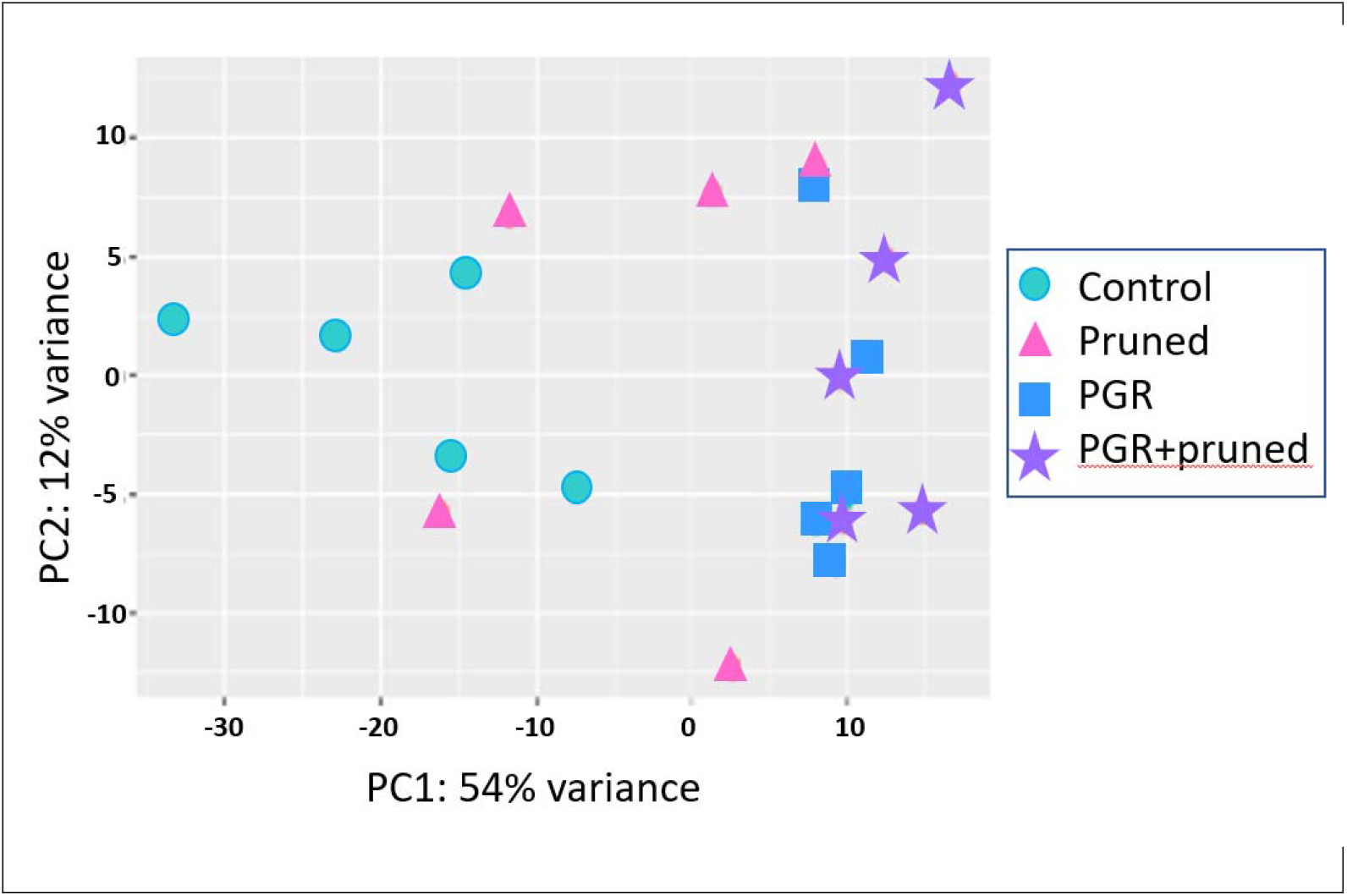
Comparison of differentially expressed genes (DEGs) by PCA with respect to treatments

**Figure 6.**
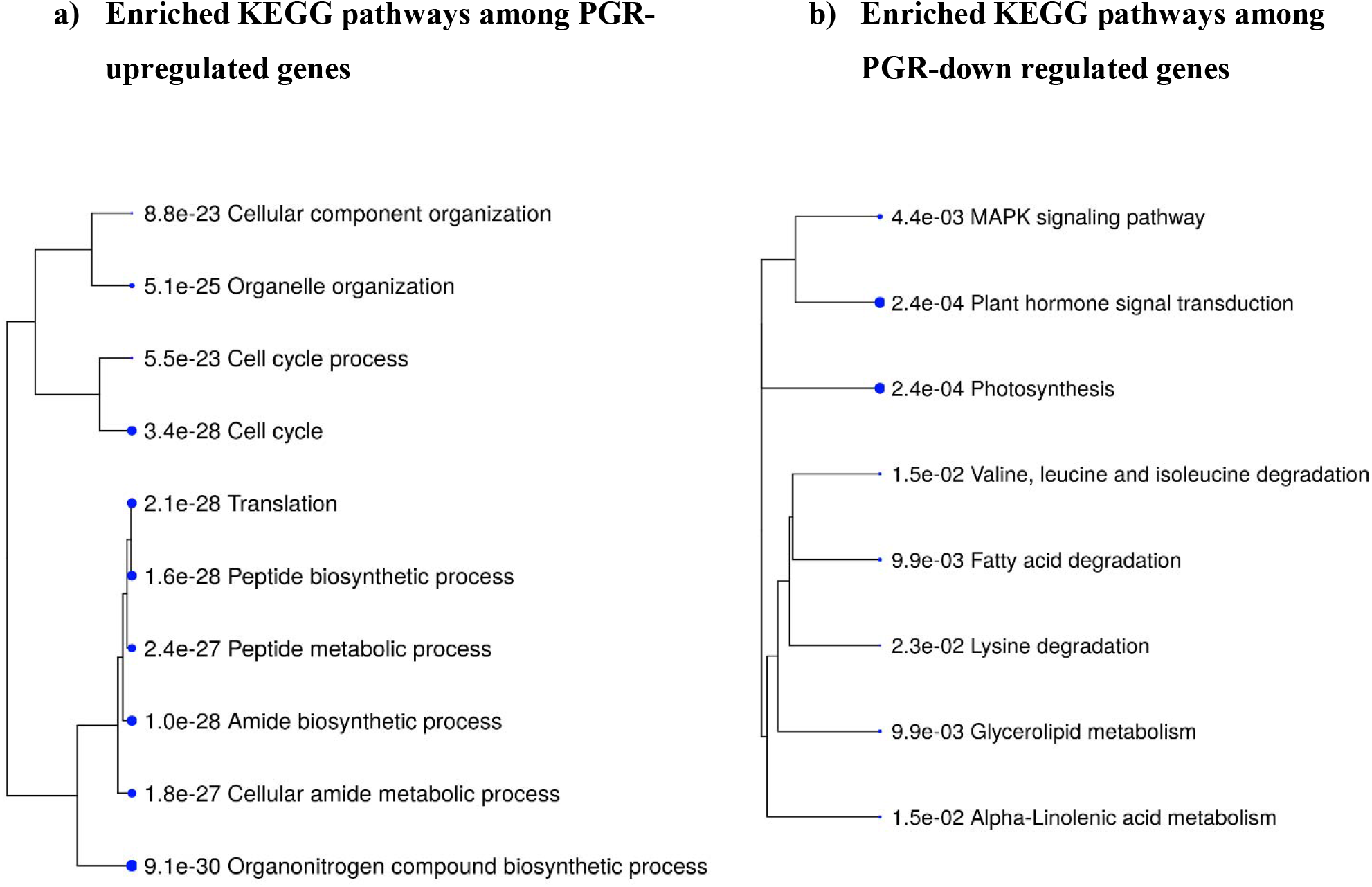
Top ten KEGG pathway enrichment terms for up-regulated and down-regulated genes in PGR-treated versus control comparison. Categories of differentially expressed genes which were significantly enriched relative to the Arabidopsis genome (p-values shown):

#### 3.3.2.1 Identification of co-expressing differentially expressed genes

Using DESeq2 statistical analysis we identified genes differentially expressed under the full model (design = ~pruning +pgr) (Figure 7a). Among the four categories of treatment (+/- PGR x +/- pruning), expression was generally grouped according to whether the category included PGR versus did not include PGR. Many genes in the plants that received the pruning treatment but were not treated with PGR were intermediate between the control and the PGR-treated plants. By inspection of the fold-expression heatmap we selected a cluster of genes with exceptionally high fold changes compared to expression profiles of other genes. Genes in this cluster had high expression in the control, very low expression in PGR-treated plants, and intermediate expression with pruning only. Enrichment analysis indicated that this cluster, consisting of 61 genes, was enriched with genes involved in abscisic acid metabolism and response, terpenoid metabolism, abiotic stress response and response to chemicals (Figure 7 b,c). This cluster suggests that regulation associated with the treatments that stimulate flowering was a decrease in stress-associated genes which were expressed at moderate levels in the control apical region.

**Figure 7.**
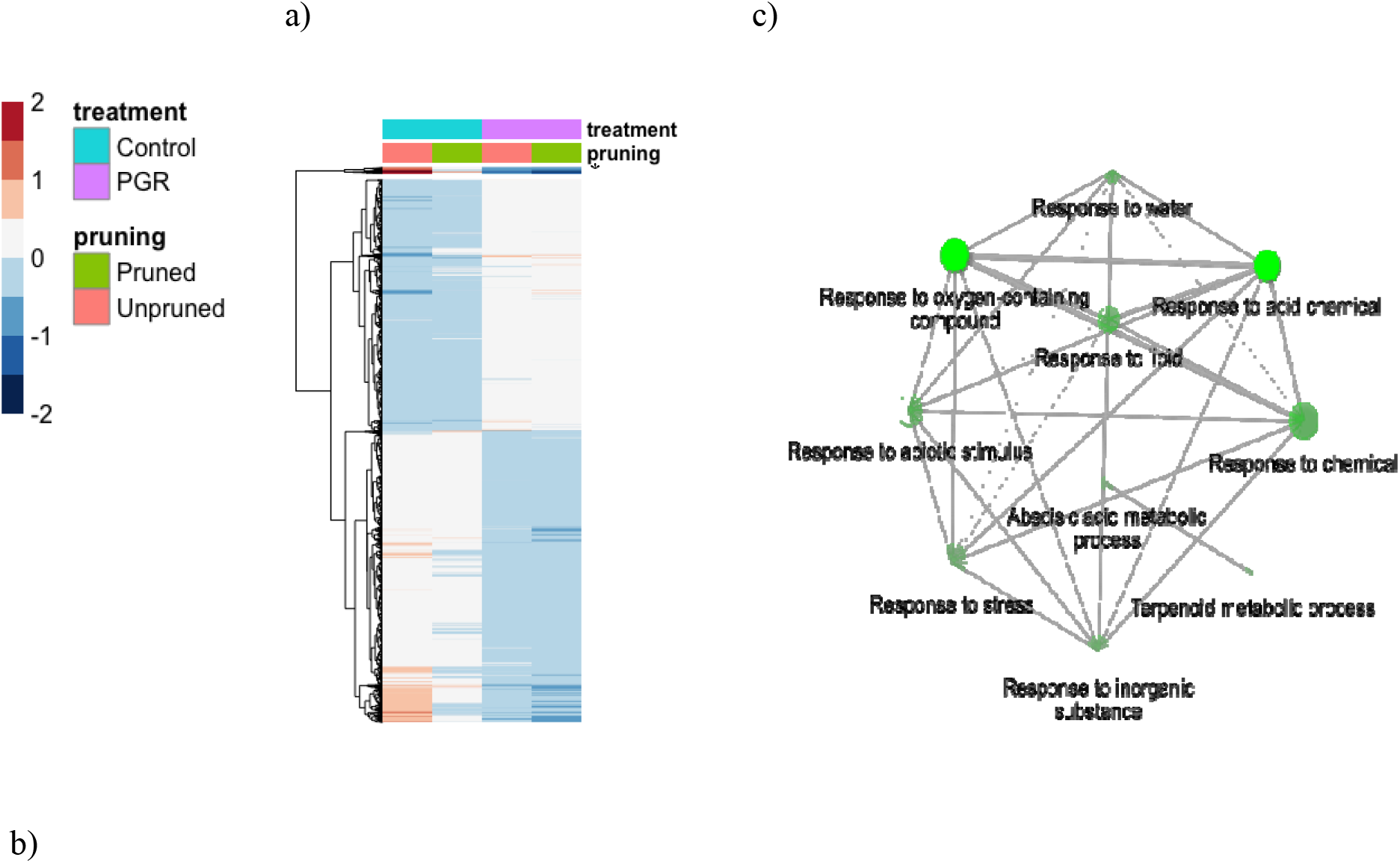
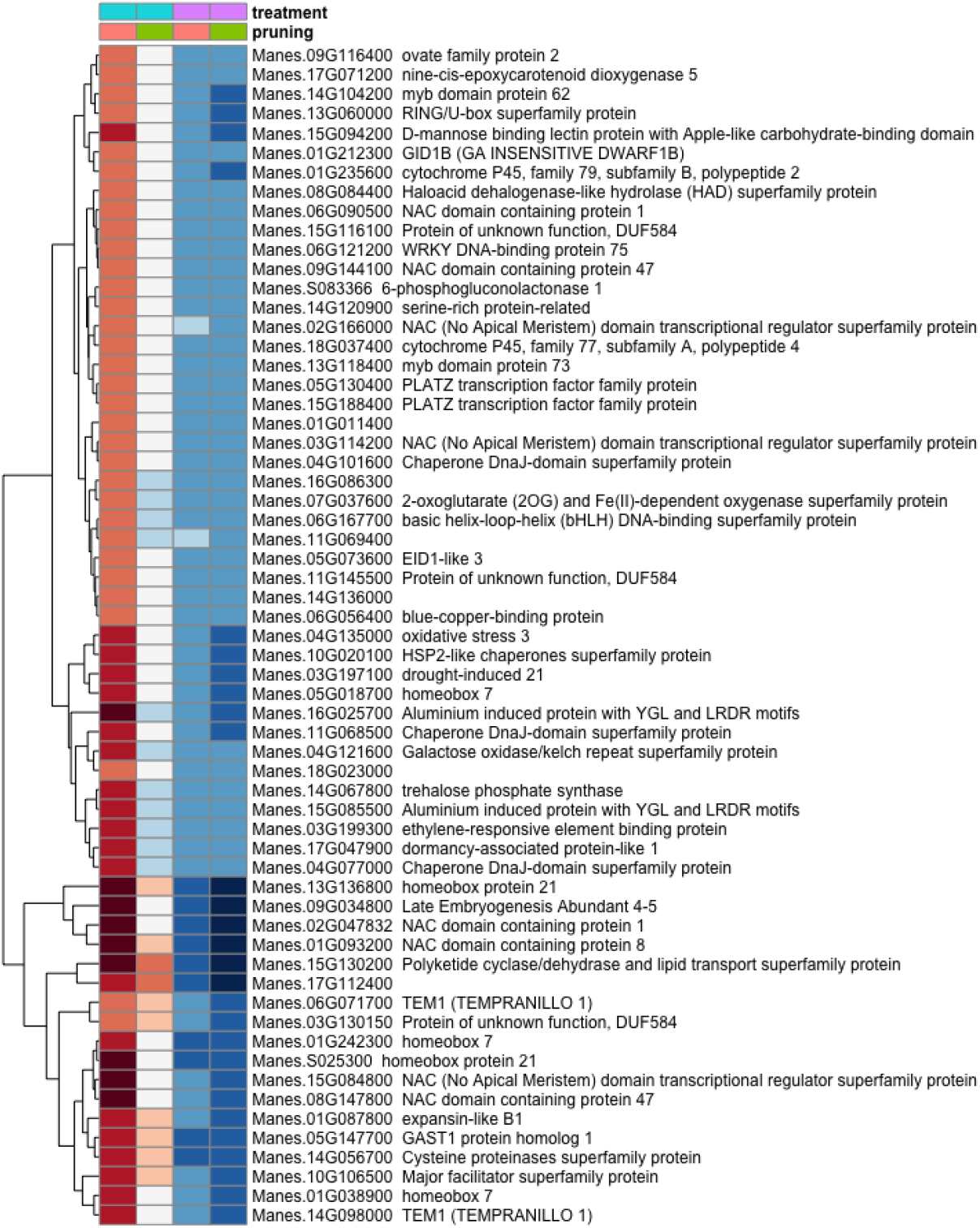
Differentially expressed genes in response to PGRs and pruning. Colors indicate fold change (log_2_ scale shown). a) Full complement of DEGs b) Uppermost slice (*) c) Enrichment analysis of the gene cluster (*)

#### 3.3.2.2 Pruning

Twenty-one genes were differentially expressed in response to pruning as the principal treatment (Figure 8a). This set was enriched in genes involved in response to wounding, herbivory and jasmonic acid signaling (Figure 8b). Also upregulated in pruning were genes involved in terpene, lipid and hormone metabolic pathways. Pruning increased expression of these genes in the presence or absence of PGR treatments. Given that the tissues for this analysis were harvested 4 d after pruning, it is not surprising that metabolic and signaling factors involved in wounding response were expressed abundantly

**Figure 8 a and b.**
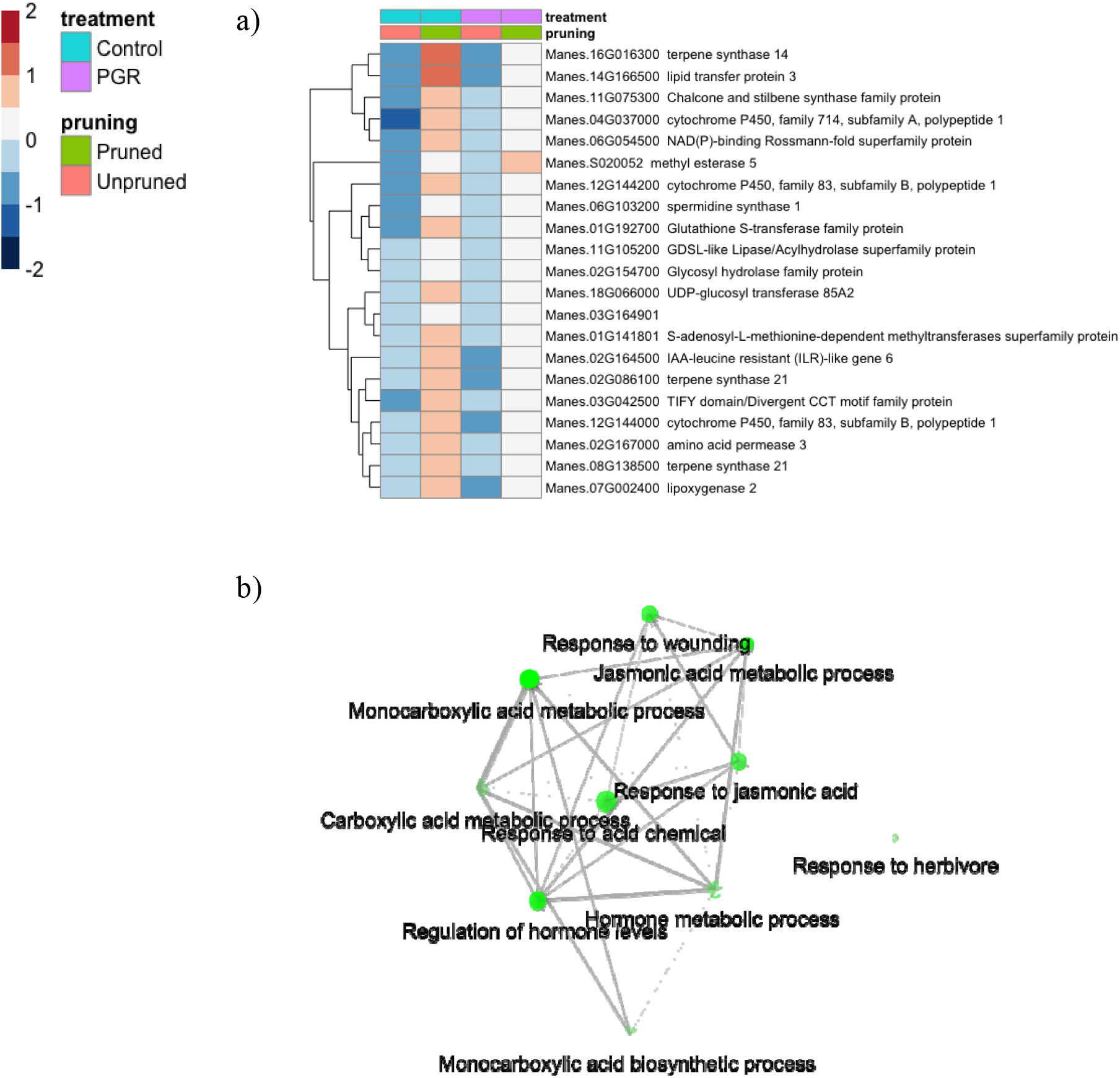
(a. top panel) differentially expressed genes identified as statistically significant in response to pruning only. (b. lower panel): Enrichment analysis of pruning responsive genes. Colors indicate fold change (log2 scale shown).

#### 3.3.2.3 Hormone signaling genes

We examined the expression profile of 115 hormone signal-transduction genes that were differentially expressed (P_adj_ < 0.05) in our samples (Figure 9; expression data for the full set of select hormones signaling genes is available in Supplementary Figure 4). While many genes differed only modestly between treatments, there was a cluster of genes with relatively high expression in the control and substantially lower expression in the PGR treatments. This cluster included a GA signaling gene (GID1b), and two repressors of ABA signaling (ABI1 and PP2CA) (Figure 9b). We also found an auxin response gene (IAA16), and a JA response gene (TIFY10B) that were expressed at a low level in the control, but they had slightly higher expression in PGR treatments, and the highest expression in the pruned without PGR treatment (Figure 9c). Thus, the expression data indicated that PGR treatments comprising anti-ethylene treatment STS and cytokinin treatment BA influenced expression of genes in hormone signaling pathways other than cytokinin and ethylene pathways as shown above, suggesting these hormones have considerable breadth of impact in the network of hormone signaling.

**Figure 9.**
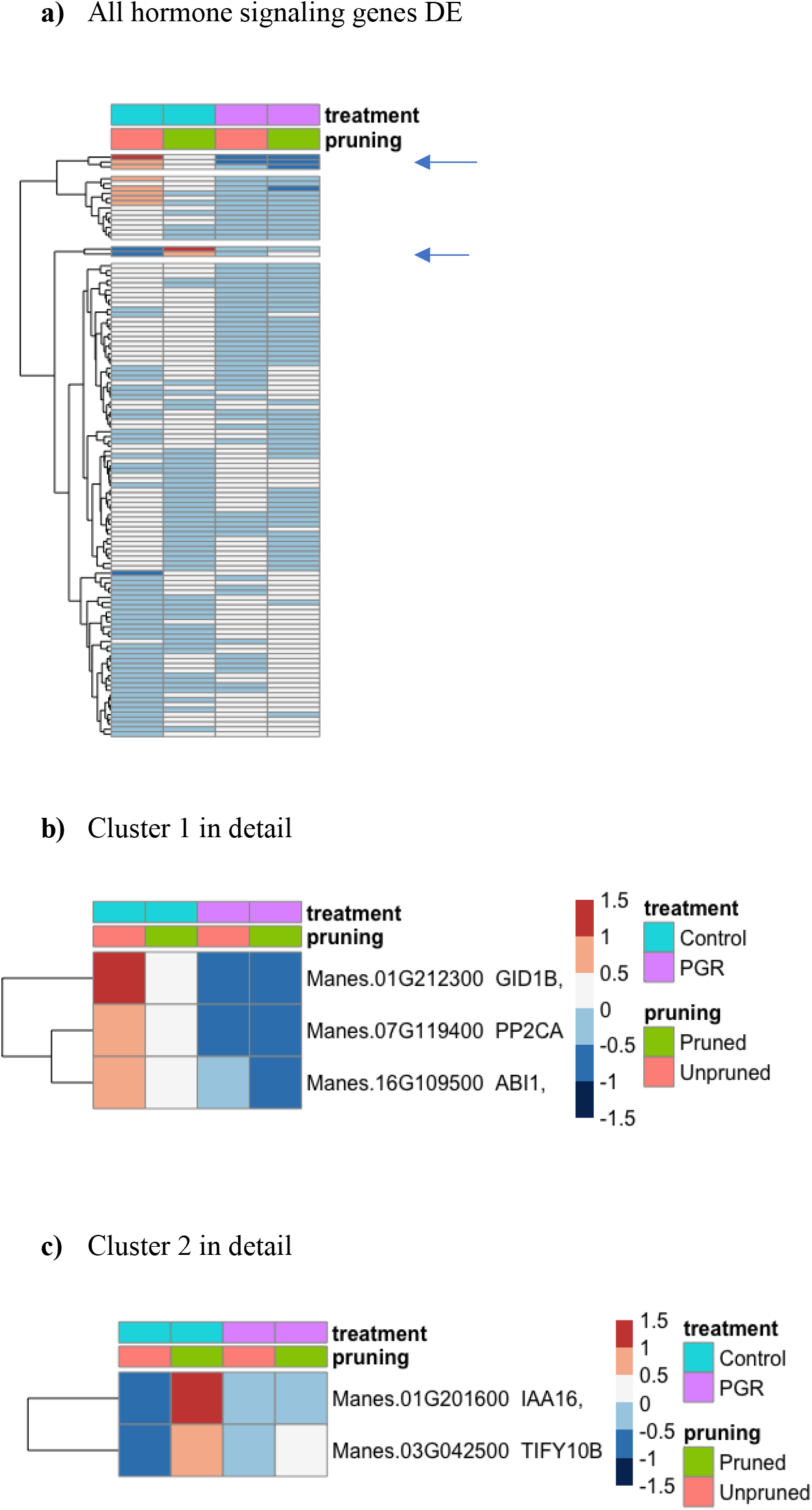
Differentially expressed hormone-signaling genes in response to PGR and pruning (a), Clusters of highly affected genes are shown in b) and c).

#### 3.3.2.4 Flowering genes

Among genes known to be involved in various aspects of flowering, we identified 217 genes that were differentially expressed (P_adj_ ≤0.05) in response to our experimental treatments (Figure 10a). From the two clusters with the largest fold changes, two members of the GA flowering pathway (GID1b, GA2ox2) and the TEM1 gene, a known flowering repressor, had the highest fold change with relatively high expression in controls, and low expression in the PGR treatments (Figure 10 b,c). Expression profiles of flowering genes organized by known flowering pathways are presented in Supplementary Figure 3. Other flowering genes with large fold changes with respect to treatments were HDA6, BRC1, FLD, NF-YA1, PNY, LUX and BRM (Figure 10 b,c).

**Figure 10.**
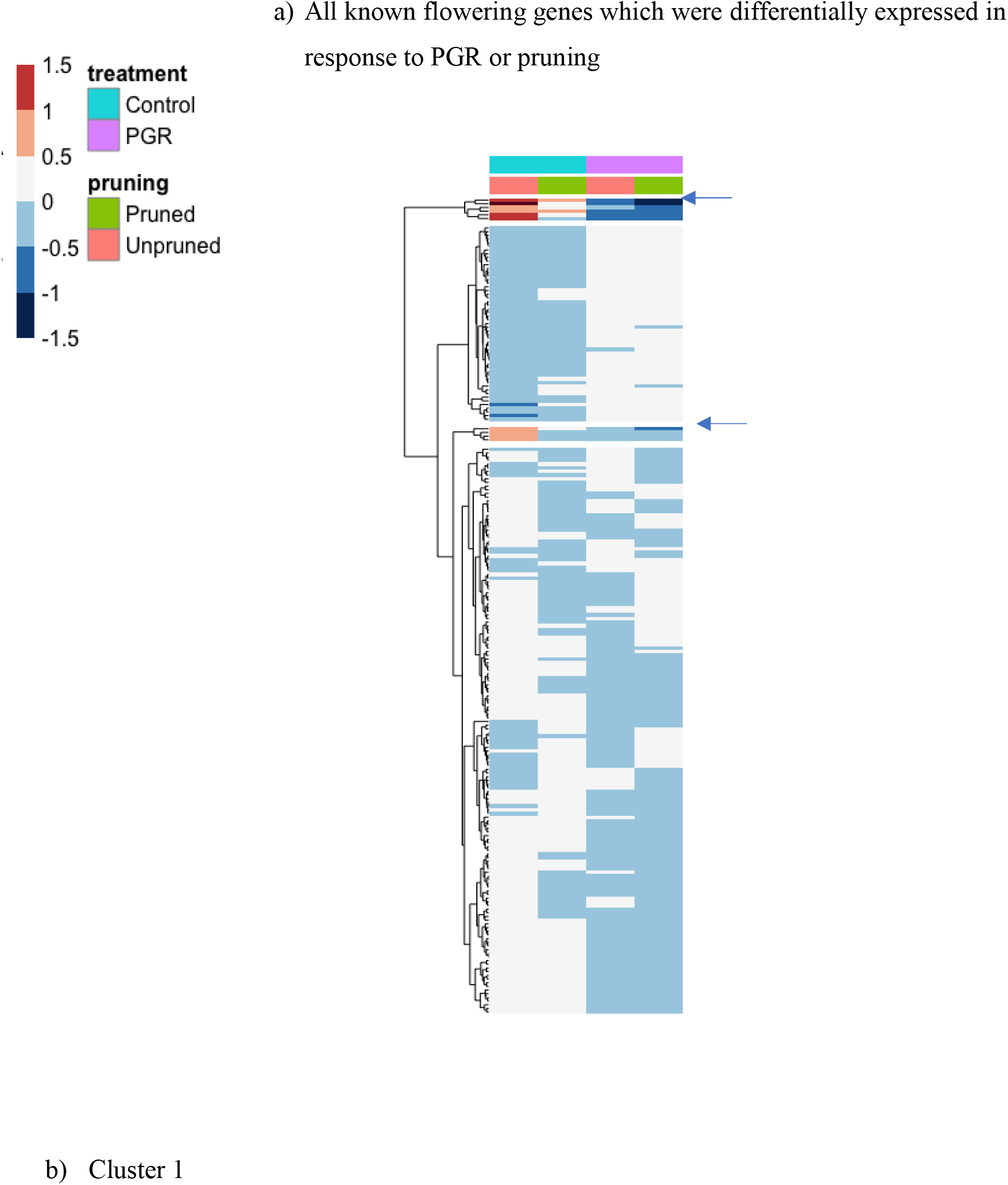
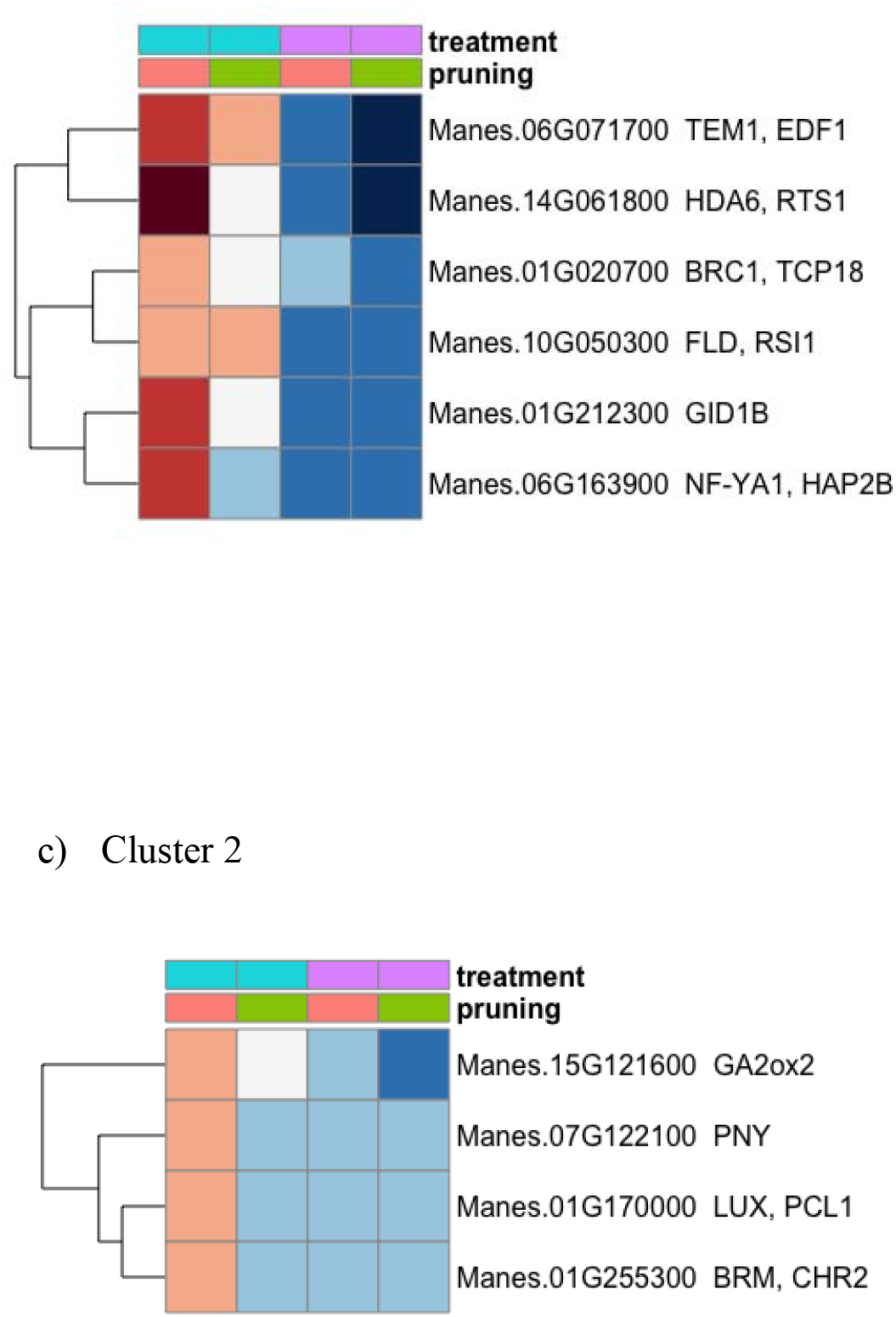
Differentially expressed flowering genes (a); details for Clusters of genes with large fold changes are shown in (b) and (c).

#### 3.3.2.5 ABCDE MADS-Box TF

Thirty of the 47 putative MADS-Box MIKC transcription factors (TFs) in cassava (http://itak.feilab.net/cgi-bin/itak/db_family_gene_list.cgi?acc=MADS-MIKC&plant=3983) were differentially expressed (P≤0.05) in response to PGR treatment or pruning (Supplementary Figure 2). The fold changes in expression levels of MADS-Box TFs were notably smaller than for the gene groups identified above. We, however, focused our attention on the flower organ development genes, with respect to the ABCDE transcription factors (Supplementary Figure 5). Generally, the A, B, C and E class genes had lower expression levels under PGR treatments while the D class genes had higher expression under PGR treatments. In contrast, an AP3 gene - Manes.02G100400 (B class) and SEP3 - Manes.13G009600 (E class) responded to pruning with increased expression with or without PGR treatment. Among these MADS-Box factors, many responded in a PGR-specific manner in which pruning without PGR was about the same as the control, but a few had lower expression in response to pruning while maintaining expression with PGR treatments (B-class AP3 Manes.02G100400, E-class SEP3 Manes.13G009600).

## 4 Discussion

### 1) STS increased total flower and fruit numbers but did not affect female to male ratio

A previous study demonstrated that STS, an anti-ethylene PGR, promotes flower development in cassava (Hyde et al. 2020),. These authors showed that STS increased the duration of cassava flower production and retention (i.e., lifespan) by five-fold relative to the control, leading to a larger number of total flowers. The present study is in line with this as STS alone increased the total number of flowers by two-fold relative to the no STS controls; STS treatment increased production of both female and male flowers, but it did not significantly alter the female to male ratios compared to the no STS control. In addition to effects on flowering, the current study also showed that STS increases fruit numbers (Figure 2, Figure 3).

In other plant systems, ethylene is widely recognized as playing a role in fruit development, especially fruit ripening (Pech et al. 2018). Ethylene hastens flower senescence, and anti-ethylene treatments including STS have been developed to increase flower longevity (Serek et al. 2006). Ethylene has also been shown to affect early stages of fruit development and fruit set. In Pea and Arabidopsis, failure to develop fruit in the absence of pollination has been associated with ovary senescence arising from increased ethylene biosynthesis in ovaries (Orzáez and Granell 1997; Carbonell-Bejerano et al. 2011). In Zucchini squash, blocking ethylene perception by STS extended ovule life span and increased the chance of developing fruit either by pollination or by parthenocarpy in response to gibberellins (Martínez et al. 2013).

In contrast to the masculinizing effect of STS on female flowers of Cannabis (Ram and Sett 1982) and Cucurbita (Den Nijs and Visser 1980), STS alone did not increase the tendency towards masculinity (Figure 2d). This could be because Cassava naturally produces a high fraction of male flowers when flowering is successful.

### 2) Pruning and STS had similar effects

Pruning of branches just below a newly initiated inflorescence has recently been shown to improve flower development in cassava and increase the total number of flowers, fruits, and seeds (Pineda et al. 2020). Hence, pruning can have similar effects to those reported for STS (Hyde et al. 2020). In addition, we observed that while both pruning and STS increased the number of total flowers, and increased female flowers as a consequence of overall increase in flower numbers, these two treatments did not affect the ratio of female to male flowers (Figure 3 and 4). The similarity in mechanisms by which STS and pruning treatments elicit their effects is open for further investigation.

### 3) BA increased female to male ratio but had no effect on total flower and fruit numbers

In other members of Euphorbiaceae, cytokinin increased the female to male ratio as illustrated in Jatropha curcas (Chen et al. 2014; Pan et al. 2014; Pan and Xu 2011) and Plukenetia volubilis (Fu et al. 2014). More female flowers were developed under higher BA concentrations (Chen et al. 2014). This was also true for cassava in our studies with BA as a sole treatment. By increasing the concentration of BA between experiments I and II, we were able to increase the percentage of female flowers from about 12% in the untreated control, to a about 30% with 0.22 mM BA to over 80% with 0.5mM BA (Figure 2 and 3). Other studies also agreed with our observations that BA was more effective at producing female flowers when treatments were begun at an early stage in flower development (Fröschle et al. 2017; Luo et al. 2020)

In contrast to the effect of BA in increasing the proportion of flowers that were female, BA had no effect on the total number of flowers and fruits. This differs from studies in other members of Euphorbiaceae in which BA treatments also resulted in an increase in the total number of flowers and fruits (Fu et al. 2014; Pan and Xu 2011; Chen et al. 2014; Pan et al. 2014). On the other hand, when either STS or pruning was combined with BA, the female to male ratio was maintained in a larger population of flowers.

### 4) Combining pruning with STS and BA has additive effects on total flower and fruit numbers while maintaining high female to male ratio

A few studies of Euphorbiaceae-family plants have shown that pruning (Pineda et al. 2020), STS (Hyde et al. 2020) or BA (Fu et al. 2014) affect flower development when applied separately, but until the current study these treatments were not applied together. Current study showed that combining these three factors substantially improved reproductive development in cassava. While STS and pruning acted additively to increase the total number of flowers, BA increased the female to male ratio of flowers. This in turn led to greater fruit development than in each of the factors applied singly (Figure 3 and 4).

### 5) Regulation of Gene expression by pruning, STS+BA and pruning combined with STS+BA

The largest fold changes in the transcriptome in response to STS+BA treatments were the down regulation of genes which were relative to their expression in the control. Among the cluster of such genes with the largest fold change (Figure 7), pathway enrichment analysis indicated that pathways related to stress were over-represented compared to their frequency in the genome. These and other examples of signaling by PGRs in our study will be further discussed below.

Genes responding to pruning as main treatment on the other hand were upregulated relative to treatments with no pruning and were enriched in processes generally related to wounding (Figure 8) (Reymond et al. 2000). This is sensible since pruning involves the excision of young fork-type branches and can thus be perceived as a type of wounding response by the plant. These genes, however, had lowered expression when pruning was combined with STS+BA, the treatment combination which produced the largest number of female flowers and fruits. This suggests that wounding related genes may not be necessary for the benefit derived from pruning in flower and fruit improvement.

### a) Hormone signaling: PGR treatments modulate GA and ABA signaling

Cassava homologues of Arabidopsis Gibberellin-Insensitive Dwarf1b (GID1b and Gibberellin 2-oxidase 2 (GA2ox2) were downregulated by PGR treatments (Figure 9 and 10). In some species, such as tea (Camellia sinensis) and Magnolia x soulangeana, GID genes are expressed at elevated levels when floral induction takes place (Jiang et al. 2020; Liu et al. 2020). GA2 oxidases on the other hand inactivate bioactive Gibberellins (GA) (Thomas et al. 1999). GA2ox2 has also been shown to be involved in the negative regulation of flowering time in combination with other GA2 oxidases (Rieu et al. 2008). The down regulation of GA2ox2 and GID1b genes by STS+BA suggest that treatments modulates GA signaling at the biosynthetic (i.e. GA2ox2) and the perception (i.e. GID1b) levels. Consistent with this role, studies of tree peony (Paeonia suffruticosa) showed that GID and GA2ox2 homologs had higher expression in the control buds than in buds stimulated to form flowers, and this response was interpreted to act in feedback regulation of GA synthesis (Guan et al. 2019).

In addition to affecting GA-related genes, PGR treatments downregulated Abscisic acid Insensitive 1 (ABI1) and Abscisic acid Hypersensitive Germination 1 (AHG1, also known as PP2CA) (Figure 9). These genes encode protein phosphatases that repress Abscisic acid (ABA) signaling and are sometimes expressed in circumstances where ABA signaling is being modulated (Lynch et al. 2012; Nishimura et al. 2007; Kuhn et al. 2006). Details into the role of ABA and the increase in sensitivity to ABA combined with a decrease in GA perception in cassava sex specification should be explored further.

### IAA16 and TIFY10b respond to pruning

The expression levels of Indoleacetic Acid-induced protein 16 (IAA16) and TIFY10b (transcriptional factors with a conserved amino acid domain - TIF[F/Y]XG) were highest in the pruning treatment, without PGRs (Figure 9). This suggests a positive relationship between these hormone-signaling genes and pruning (Korasick et al. 2014). The similar expression patterns of IAA16 and TIFY10b suggest a possible role in cassava pruning response which should be probed further.

### b) PGR and pruning treatments modulate flowering genes

In the current study, a group of flowering related genes were downregulated by STS+BA treatment (Figure 10). These included the positive flowering effectors Histone Deacetylase 6 (HDA6), and NUCLEAR FACTOR Y, SUBUNIT A1(NF–YA1), and negative flowering effectors Branched 1 (BRC1), and Brahma1 (BRM1, an ATP dependent chromatin remodeler). In addition to their roles in flowering (Bouché et al. 2016), these genes also possess functions related to ABA response (Chen et al. 2010; Peirats-Llobet et al. 2016; González-Grandío et al. 2017; Mu et al. 2013). These all had similar expression patterns: they were strongly down regulated by PGR treatment. BRM1 was an exception to this as it was only moderately downregulated by PGR treatment.

Among the strongly downregulated genes, the floral repressor Tempranillo 1 (TEM1), functions as a downstream effector of ethylene responses and also suppresses the biosynthesis of bioactive GA (Osnato et al. 2012). The downregulation of TEM1 by floral enhancing treatments in this study, may reflect a role for it beyond its association with juvenility (Sgamma et al. 2013) and role as a flowering repressor, possibly providing a link between repressed GA and ethylene signaling (Matias-Hernandez 2014). This is reasonable given that the mode of action of STS is silver ion binding to the ethylene receptor and thereby inhibiting ethylene perception (Veen 1983).

## 3.5 Conclusion

This study showed that combining pruning, with anti-ethylene STS (supplied by petiole feeding) and cytokinin BA (sprayed to the shoot apex) significantly improved cassava female flower and fruit development. It also demonstrated that female flower enhancing treatments led to transcriptional changes indicating that the treatments repressed ethylene and GA signaling and modulated ABA signaling. Our work contributes to understanding of sex determination and to a body of evidence on hormone and signaling factors in floral development. This work also provides a flowering protocol that will facilitate cassava breeding and contribute to global food security.

## 6 Author Contributions

D.O, P.K and T.S obtained funding. D.O and T.S designed experiment. P.K supervised work on field in Nigeria. D.O, O.E and P.H conducted field and controlled condition experiments. D.O analysed data. D.O, O.E, P.H, P.K, T.S wrote manuscript.

## 8 Acknowledgments

The authors would also like to thank the Federal Government of Nigeria for partly funding Deborah’s studies through the Presidential Special Scholarship for Innovation and Development (PRESSID) managed by the National Universities Commission (NUC) and funded by the Federal Scholarship Board (FSB). The authors would also like to thank the Bioscience unit and Cassava Breeding unit of the International Institute of Tropical Agriculture, Nigeria for providing laboratory facilities, office space and field space for conducting experiments.

This research was part of the Ph.D. dissertation of the first author at Cornell University, Ithaca, NY

## References

Adeyemo OS, Chavarriaga P, Tohme J, Fregene M, Davis SJ, Setter TL (2017) Overexpression of Arabidopsis FLOWERING LOCUS T (FT) gene improves floral development in cassava (Manihot esculenta, Crantz). Plos One 12 (7):e0181460

Adeyemo OS, Hyde PT, Setter TL (2019) Identification of FT family genes that respond to photoperiod, temperature and genotype in relation to flowering in cassava (Manihot esculenta, Crantz). Plant reproduction 32 (2):181–191

An J, Almasaud RA, Bouzayen M, Zouine M, Chervin C (2020) Auxin and ethylene regulation of fruit set. Plant Sci 292:110381

Bouché F, Lobet G, Tocquin P, Périlleux C (2016) FLOR-ID: an interactive database of flowering-time gene networks in Arabidopsis thaliana. Nucleic Acids Research 44 (D1):D1167–D1171

Brooks ME, Kristensen K, van Benthem KJ, Magnusson A, Berg CW, Nielsen A, Skaug HJ, Machler M, Bolker BM (2017) glmmTMB balances speed and flexibility among packages for zero-inflated generalized linear mixed modeling. The R journal 9 (2):378–400

Bull SE, Alder A, Barsan C, Kohler M, Hennig L, Gruissem W, Vanderschuren H (2017) FLOWERING LOCUS T triggers early and fertile flowering in glasshouse cassava (Manihot esculenta Crantz). Plants 6 (2):22

Carbonell-Bejerano P, Urbez C, Granell A, Carbonell J, Perez-Amador MA (2011) Ethylene is involved in pistil fate by modulating the onset of ovule senescence and the GA-mediated fruit set in Arabidopsis. Bmc Plant Biol 11 (1):84

Ceballos H, Iglesias CA, Perez JC, Dixon AG (2004) Cassava breeding: opportunities and challenges. Plant Mol Biol 56 (4):503–516

Ceballos H, Pérez JC, Joaqui Barandica O, Lenis JI, Morante N, Calle F, Pino L, Hershey CH (2016) Cassava breeding I: The value of breeding value. Front Plant Sci 7:1227

Chen L-T, Luo M, Wang Y-Y, Wu K (2010) Involvement of Arabidopsi s histone deacetylase HDA6 in ABA and salt stress response. J Exp Bot 61 (12):3345–3353

Chen M-S, Pan B-Z, Wang G-J, Ni J, Niu L, Xu Z-F (2014) Analysis of the transcriptional responses in inflorescence buds of Jatropha curcasexposed to cytokinin treatment. Bmc Plant Biol 14 (1):318

De Souza AP, Long SP (2018) Toward improving photosynthesis in cassava: characterizing photosynthetic limitations in four current African cultivars. Food and energy security 7 (2):e00130

Den Nijs A, Visser D (1980) Induction of male flowering in gynoecious cucumbers (Cucumis sativus L.) by silver ions. Euphytica 29 (2):273–280

Dennis G, Sherman BT, Hosack DA, Yang J, Gao W, Lane HC, Lempicki RA (2003) DAVID: database for annotation, visualization, and integrated discovery. Genome biology 4 (9):1–11

Fröschle M, Horn H, Spring O (2017) Effects of the cytokinins 6-benzyladenine and forchlorfenuron on fruit-, seed-and yield parameters according to developmental stages of flowers of the biofuel plant Jatropha curcas L.(Euphorbiaceae). Plant Growth Regul 81 (2):293–303

Fu Q, Niu L, Zhang Q, Pan B-Z, He H, Xu Z-F (2014) Benzyladenine treatment promotes floral feminization and fruiting in a promising oilseed crop Plukenetia volubilis. Industrial Crops and Products 59:295–298

Ge SX, Jung D, Yao R (2020) ShinyGO: a graphical gene-set enrichment tool for animals and plants. Bioinformatics 36 (8):2628–2629

Giovannoni JJ (2004) Genetic regulation of fruit development and ripening. The plant cell 16 (suppl 1):S170–S180

González-Grandío E, Pajoro A, Franco-Zorrilla JM, Tarancón C, Immink RG, Cubas P (2017) Abscisic acid signaling is controlled by a BRANCHED1/HD-ZIP I cascade in Arabidopsis axillary buds. Proceedings of the National Academy of Sciences 114 (2):E245–E254

Goodstein DM, Shu S, Howson R, Neupane R, Hayes RD, Fazo J, Mitros T, Dirks W, Hellsten U, Putnam N (2012) Phytozome: a comparative platform for green plant genomics. Nucleic acids research 40 (D1):D1178–D1186

Guan Y.R, Xue J.Q, Xue Y.Q, Yang R.W, Wang S.L, Zhang X.X (2019) Effect of exogenous GA3 on flowering quality, endogenous hormones, and hormone-and flowering-associated gene expression in forcing-cultured tree peony (Paeonia suffruticosa). Journal of Integrative Agriculture 18 (6):1295–1311

Halsey ME, Olsen KM, Taylor NJ, Chavarriaga Aguirre P (2008) Reproductive biology of cassava (Manihot esculenta Crantz) and isolation of experimental field trials. Crop Sci 48 (1):49–58

Horn JW, van Ee BW, Morawetz JJ, Riina R, Steinmann VW, Berry PE, Wurdack KJ (2012) Phylogenetics and the evolution of major structural characters in the giant genus Euphorbia L.(Euphorbiaceae). Molecular Phylogenetics and Evolution 63 (2):305–326

Hyde PT, Guan X, Abreu V, Setter TL (2020) The anti-ethylene growth regulator silver thiosulfate (STS) increases flower production and longevity in cassava (Manihot esculenta Crantz). Plant Growth Regul 90 (3):441–453

Iragaba P (2019) Translating Cassava Attributes Preferred by Ugandan Smallholder Farmers into Breeding Targets. Cornell University,

Jiang Z, Sun L, Wei Q, Ju Y, Zou X, Wan X, Liu X, Yin Z (2020) A New Insight into Flowering Regulation: Molecular Basis of Flowering Initiation in Magnolia× soulangeana ‘Changchun’. Genes 11 (1):15

Kanrar S, Bhattacharya M, Arthur B, Courtier J, Smith HM (2008) Regulatory networks that function to specify flower meristems require the function of homeobox genes PENNYWISE and POUND□FOOLISH in Arabidopsis. The Plant Journal 54 (5):924–937

Korasick DA, Westfall CS, Lee SG, Nanao MH, Dumas R, Hagen G, Guilfoyle TJ, Jez JM, Strader LC (2014) Molecular basis for AUXIN RESPONSE FACTOR protein interaction and the control of auxin response repression. Proceedings of the National Academy of Sciences 111 (14):5427–5432

Kuhn JM, Boisson-Dernier A, Dizon MB, Maktabi MH, Schroeder JI (2006) The protein phosphatase AtPP2CA negatively regulates abscisic acid signal transduction in Arabidopsis, and effects of abh1 on AtPP2CA mRNA. Plant Physiol 140 (1):127–139

Lenth R (2019) emmeans: Estimated marginal means, aka least-squares means. R package version 1.4. 3.01.

Li S, Cui Y, Zhou Y, Luo Z, Liu J, Zhao M (2017) The industrial applications of cassava: current status, opportunities and prospects. Journal of the Science of Food and Agriculture 97 (8):2282–2290

Lin Y-H, Lin M-H, Gresshoff PM, Ferguson BJ (2011) An efficient petiole-feeding bioassay for introducing aqueous solutions into dicotyledonous plants. Nature Protocols 6 (1):36–45

Liu Q, Liu J, Zhang P, He S (2014) Root and tuber crops.

Liu X, Hou X (2018) Antagonistic regulation of ABA and GA in metabolism and signaling pathways. Front Plant Sci 9:251

Liu Y, Hao X, Lu Q, Zhang W, Zhang H, Wang L, Yang Y, Xiao B, Wang X (2020) Genome-wide identification and expression analysis of flowering-related genes reveal putative floral induction and differentiation mechanisms in tea plant (Camellia sinensis). Genomics

Love MI, Huber W, Anders S (2014) Moderated estimation of fold change and dispersion for RNA-seq data with DESeq2. Genome biology 15 (12):550

Luo Y, Pan B-Z, Li L, Yang C-X, Xu Z-F (2020) Developmental basis for flower sex determination and effects of cytokinin on sex determination in Plukenetia volubilis (Euphorbiaceae). Plant Reproduction:1–14

Lynch T, Erickson BJ, Finkelstein RR (2012) Direct interactions of ABA-insensitive (ABI)-clade protein phosphatase (PP) 2Cs with calcium-dependent protein kinases and ABA response element-binding bZIPs may contribute to turning off ABA response. Plant Mol Biol 80 (6):647–658

Martínez C, Manzano S, Megías Z, Garrido D, Picó B, Jamilena M (2013) Involvement of ethylene biosynthesis and signalling in fruit set and early fruit development in zucchini squash (Cucurbita pepo L.). Bmc Plant Biol 13 (1):139

Matias-Hernandez L. 2014. RAV genes: regulation of floral induction and beyond. Annals of Botany (London) 114, 1459–1470.

Mishra NS, Tuteja R, Tuteja N (2006) Signaling through MAP kinase networks in plants. Archives of Biochemistry and Biophysics 452 (1):55–68

Moormann F, Lal R, Juo A (1975) The soils of IITA, IITA Technical Bulletin No. 3. IITA, Ibadan

Mu J, Tan H, Hong S, Liang Y, Zuo J (2013) Arabidopsis transcription factor genes NF-YA1, 5, 6, and 9 play redundant roles in male gametogenesis, embryogenesis, and seed development. Mol Plant 6 (1):188–201

Nassar NM (1980) Attempts to hybridize wildManihot species with cassava. Economic Botany 34 (1):13–15

Nishimura N, Yoshida T, Kitahata N, Asami T, Shinozaki K, Hirayama T (2007) ABA□Hypersensitive Germination1 encodes a protein phosphatase 2C, an essential component of abscisic acid signaling in Arabidopsis seed. The Plant Journal 50 (6):935–949

Odipio J, Getu B, Chauhan R, Alicai T, Bart R, Nusinow DA, Taylor NJ (2020) Transgenic overexpression of endogenous FLOWERING LOCUS T-like gene MeFT1 produces early flowering in cassava. Plos One 15 (1):e0227199

Onai K, Ishiura M (2005) PHYTOCLOCK 1 encoding a novel GARP protein essential for the Arabidopsis circadian clock. Genes to Cells 10 (10):963–972

Orzáez D, Granell A (1997) DNA fragmentation is regulated by ethylene during carpel senescence in Pisum sativum. The Plant Journal 11 (1):137–144

Osnato M, Castillejo C, Matías-Hernández L, Pelaz S (2012) TEMPRANILLO genes link photoperiod and gibberellin pathways to control flowering in Arabidopsis. Nat Commun 3 (1):1–8

Pan B-Z, Chen M-S, Ni J, Xu Z-F (2014) Transcriptome of the inflorescence meristems of the biofuel plant Jatropha curcas treated with cytokinin. BMC genomics 15 (1):974

Pan B-Z, Xu Z-F (2011) Benzyladenine treatment significantly increases the seed yield of the biofuel plant Jatropha curcas. J Plant Growth Regul 30 (2):166–174

Pech JC, Purgatto E, Bouzayen M, Latché A (2018) Ethylene and fruit ripening. Annual Plant Reviews online:275–304

Peirats-Llobet M, Han S-K, Gonzalez-Guzman M, Jeong CW, Rodriguez L, Belda-Palazon B, Wagner D, Rodriguez PL (2016) A direct link between abscisic acid sensing and the chromatin-remodeling ATPase BRAHMA via core ABA signaling pathway components. Mol Plant 9 (1):136–147

Perera PI, Quintero M, Dedicova B, Kularatne J, Ceballos H (2013) Comparative morphology, biology and histology of reproductive development in three lines of Manihot esculenta Crantz (Euphorbiaceae: Crotonoideae). Aob Plants 5

Pineda LM, Yu B, Tian Y, Morante N, Salazar S, Hyde P, Setter TL, Ceballos H. 2020. Effect of pruning young branches on fruit and seed set in cassava. Frontiers in Plant Science 11, 1107.

Ram HM, Sett R (1982) Induction of fertile male flowers in genetically female Cannabis sativa plants by silver nitrate and silver thiosulphate anionic complex. Theor Appl Genet 62 (4):369–375

Reymond P, Weber H, Damond M, Farmer EE. 2000. Differential Gene Expression in Response to Mechanical Wounding and Insect Feeding in Arabidopsis Plant Cell 12, 707–720.

Rieu I, Eriksson S, Powers SJ, Gong F, Griffiths J, Woolley L, Benlloch R, Nilsson O, Thomas SG, Hedden P (2008) Genetic analysis reveals that C19-GA 2-oxidation is a major gibberellin inactivation pathway in Arabidopsis. The Plant Cell 20 (9):2420–2436

Rife TW, Poland JA (2014) Field Book: An open□source application for field data collection on Android. Crop Science 54 (4):1624–1627

Serek M, Woltering EJ, Sisler EC, Frello S, Sriskandarajah S. 2006. Controlling ethylene responses in flowers at the receptor level. Biotechnology Advances 24, 368–381.

Sgamma T, Jackson A, Muleo R, Thomas B, Massiah A. 2014. TEMPRANILLO is a regulator of juvenility in plants. Scientific Reports 4, 3704.

Sonnewald U, Fernie AR, Gruissem W, Schläpfer P, Anjanappa RB, Chang SH, Ludewig F, Rascher U, Muller O, van Doorn AM (2020) The Cassava Source Sink project: Opportunities and challenges for crop improvement by metabolic engineering. The Plant Journal

Souza LS, Alves AAC, de Oliveira EJ (2020) Phenological diversity of flowering and fruiting in cassava germplasm. Sci Hortic-Amsterdam 265:109253

Team RC (2013) R: A language and environment for statistical computing.

Theißen G, Melzer R, Rümpler F (2016) MADS-domain transcription factors and the floral quartet model of flower development: linking plant development and evolution. Development 143 (18):3259–3271

Thomas SG, Phillips AL, Hedden P (1999) Molecular cloning and functional expression of gibberellin 2-oxidases, multifunctional enzymes involved in gibberellin deactivation. Proceedings of the National Academy of Sciences 96 (8):4698–4703

Tuteja N, Gill SS, Tiburcio AF, Tuteja R (2012) Improving crop resistance to abiotic stress. John Wiley & Sons,

Vanholme B, Grunewald W, Bateman A, Kohchi T, Gheysen G (2007) The tify family previously known as ZIM. Trends Plant Sci 12 (6):239–244

Veen H (1983) Silver thiosulphate: an experimental tool in plant science. Sci Hortic-Amsterdam 20 (3):211–224

Wolfe MD, Del Carpio DP, Alabi O, Ezenwaka LC, Ikeogu UN, Kayondo IS, Lozano R, Okeke UG, Ozimati AA, Williams E (2017) Prospects for genomic selection in cassava breeding. The Plant Genome 10 (3):1–19

Yang C-H, Chou M-L (1999) FLD interacts with CO to affect both flowering time and floral initiation in Arabidopsis thaliana. Plant Cell Physiol 40 (6):647–650

Zheng Y, Jiao C, Sun H, Rosli HG, Pombo MA, Zhang P, Banf M, Dai X, Martin GB, Giovannoni JJ (2016) iTAK: a program for genome-wide prediction and classification of plant transcription factors, transcriptional regulators, and protein kinases. Mol Plant 9 (12):1667–1670

