## Supplementary data for "Silver thiosulfate and Benzyladenine in combination with pruning additively feminizes cassava flowers and modulates transcriptome"

***Supplementary Material***

| a) IITA-TMS-IBA980002 | |
| --- | --- |
| 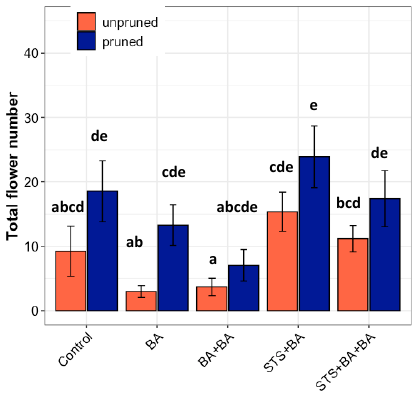 | 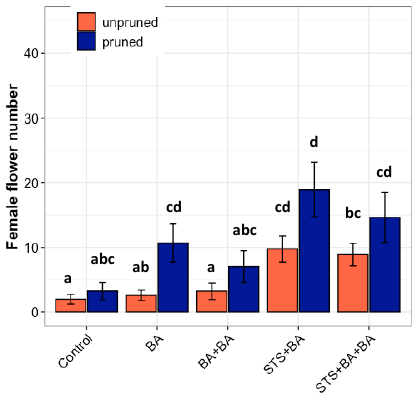 |
| 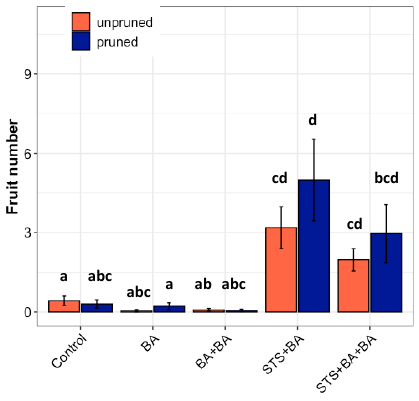 | 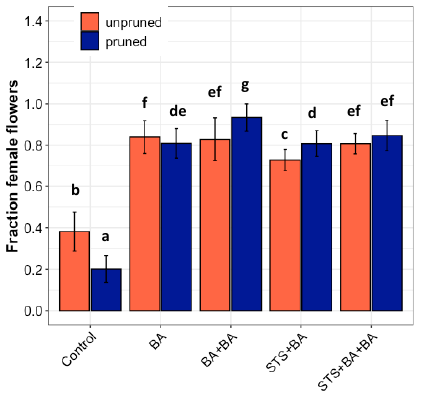 |

| b) IITA-TMS-IBA30572 | |
| --- | --- |
| 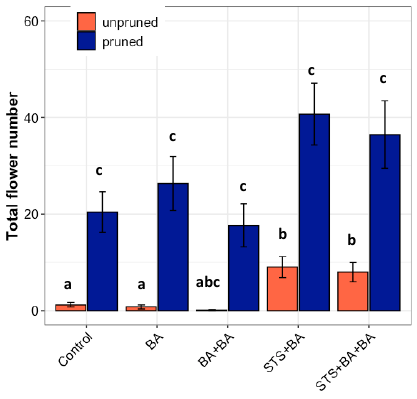 | 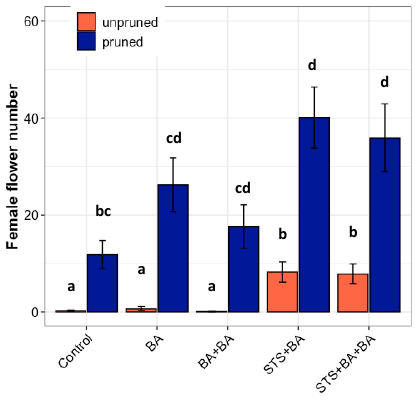 |
| 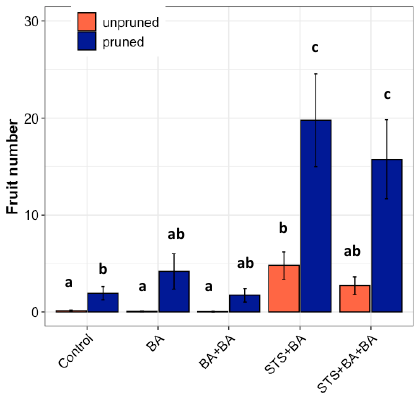 | 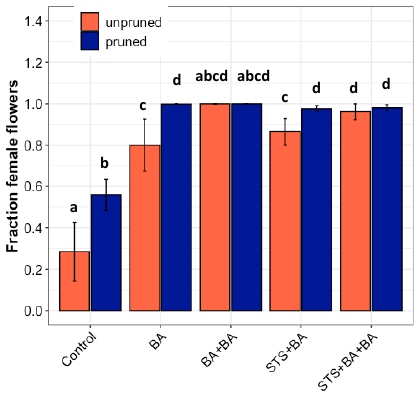 |

| c) TMEB419 | |
| --- | --- |
| 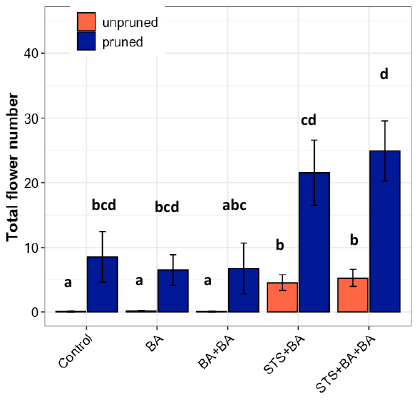 | 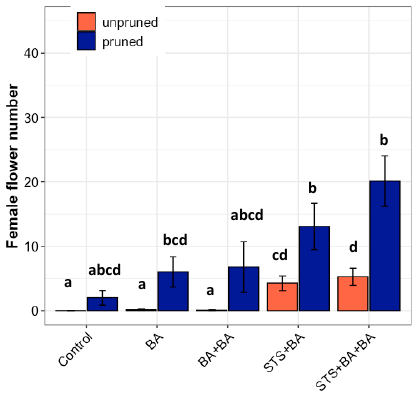 |
| 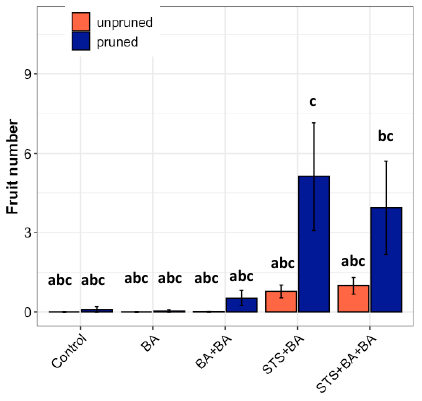 | 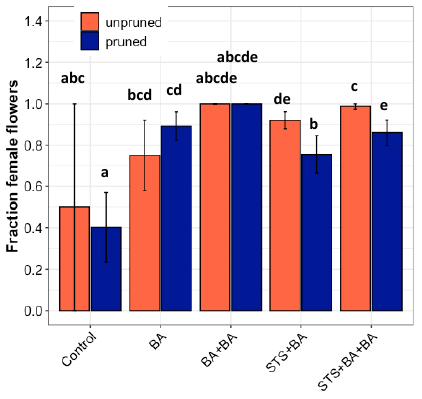 |

**Supplementary Figure 1** Flowering response to treatment by genotypes a) IITA-TMS-IBA980002 b) IITA-TMS-IBA30572 c) TMEB419

Scale

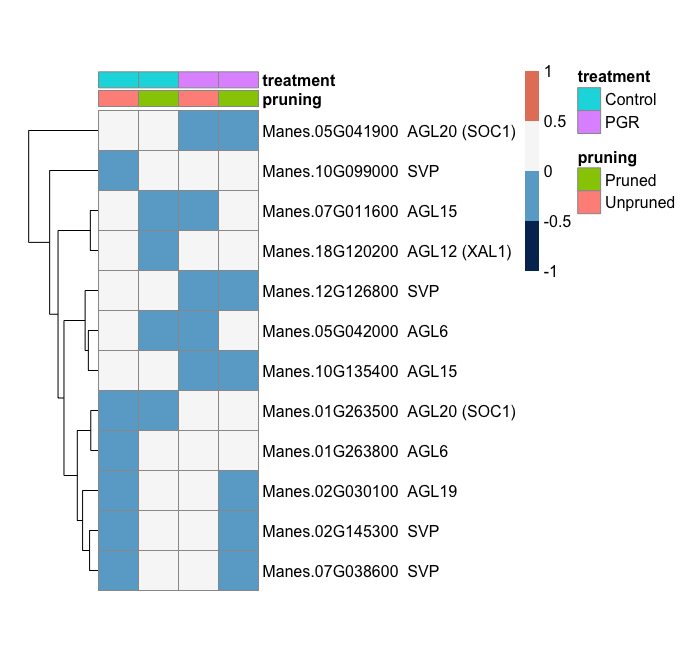

a) All MADS-Box genes

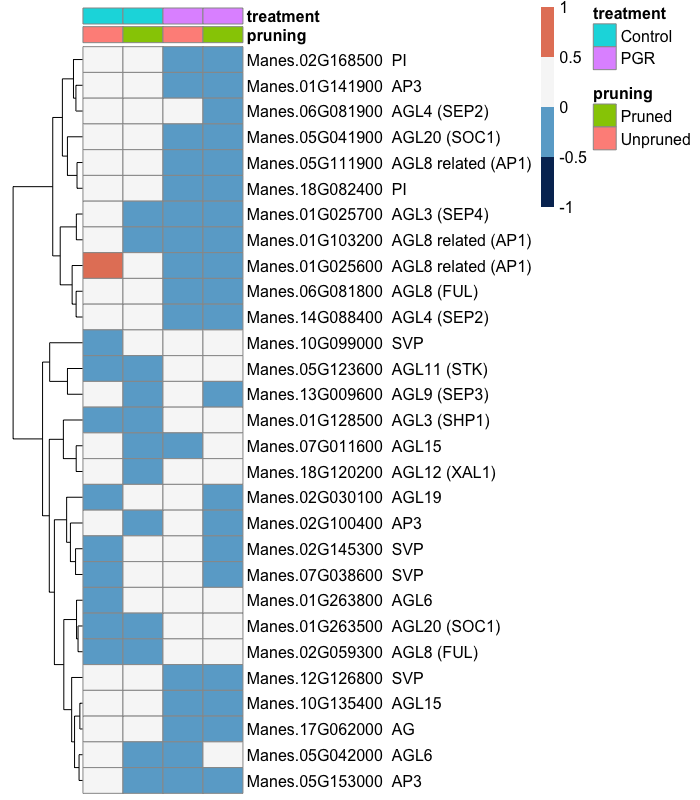

b) Floral transition MADSbox genes

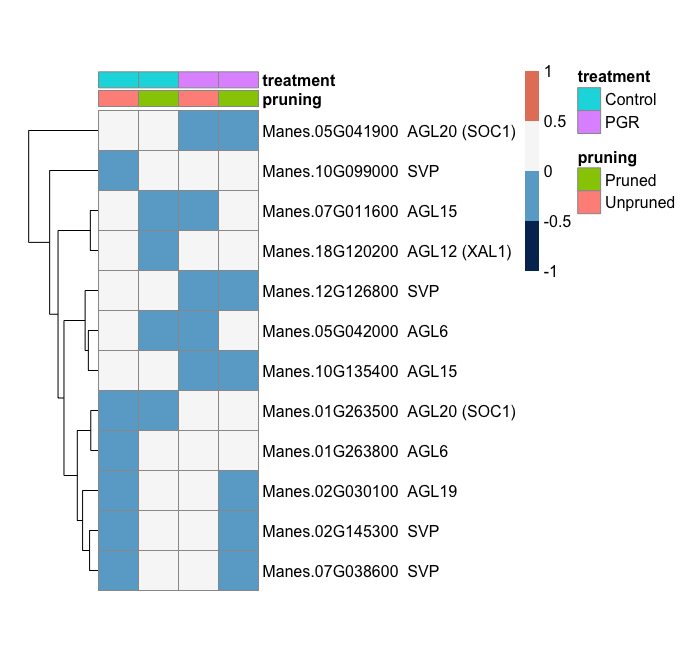

c) Floral and inflorescence MADSBox meristem genes

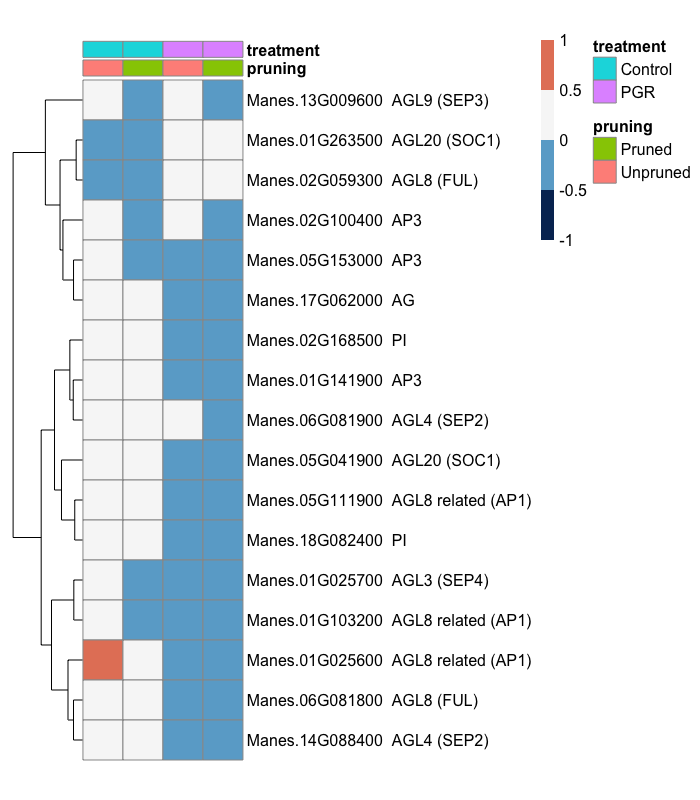

**Supplementary Figure 2** All MADS Box MIKC gene differentially expressed .a) All MADS-Box genes b) Floral transition MADSbox genes c) Floral and inflorescence MADSBox meristem genes

Scale

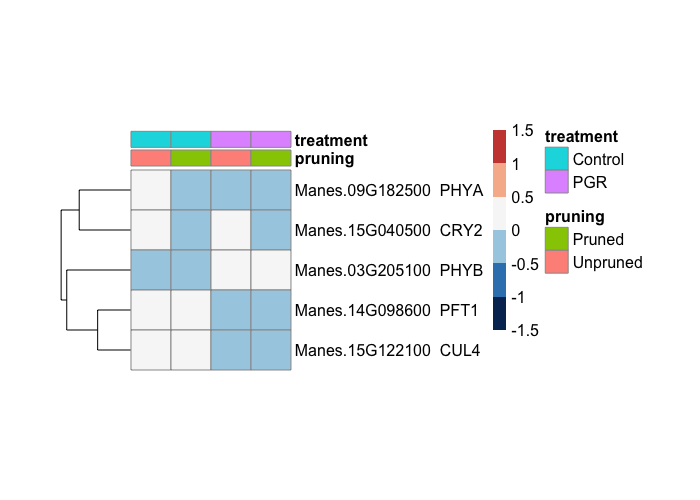

a) Flowering genes: vegetative to reproductive phase change

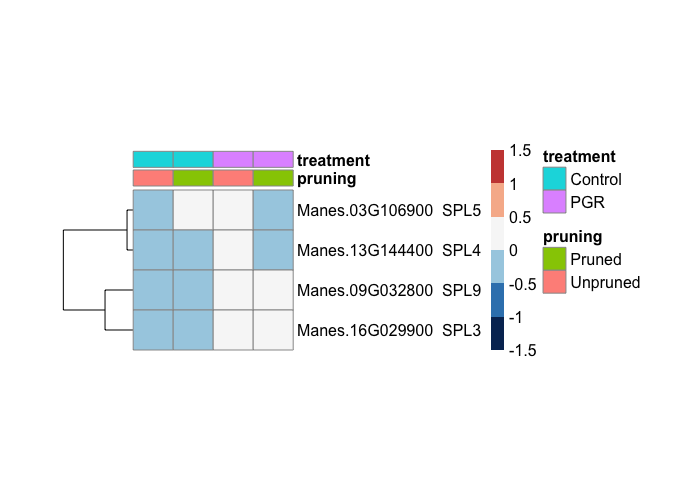

b) Flowering genes: Light perception

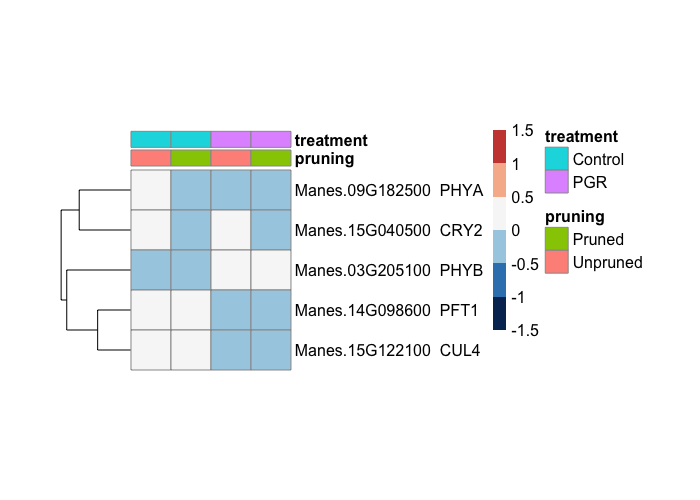

c) Flowering genes: GA pathway

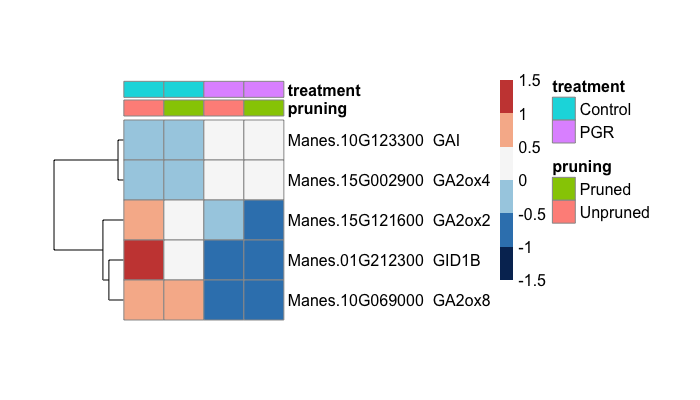

d) Flowering genes: repressor

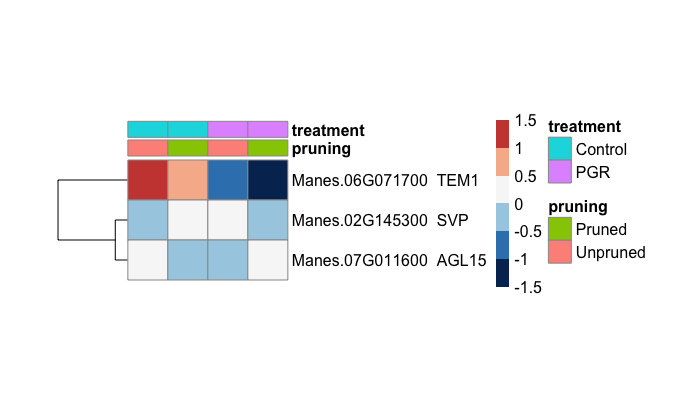

e) Flowering genes: meristem genes

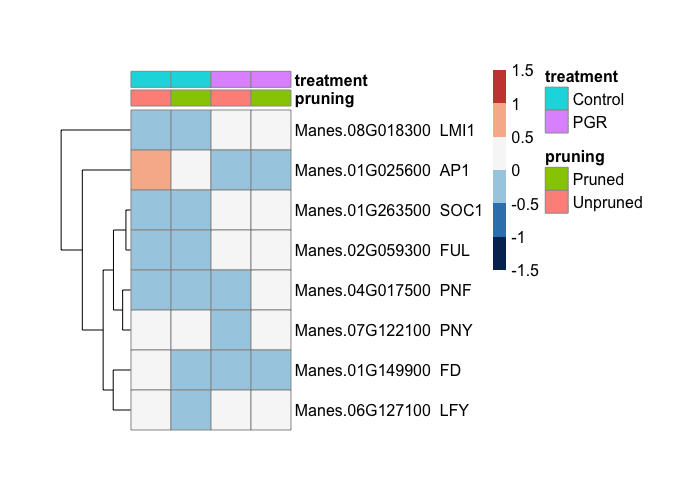

f) Flowering genes: Vernalization

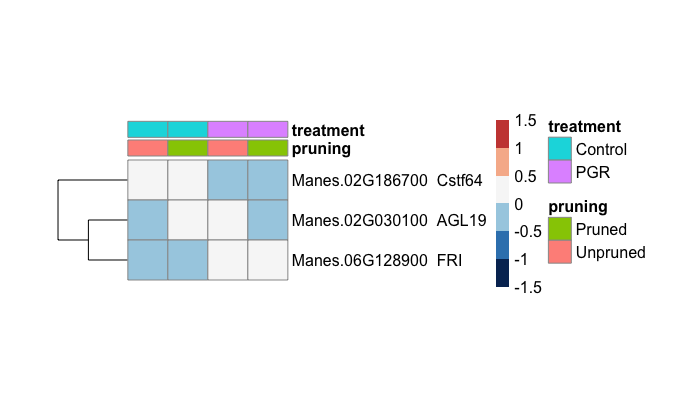

g) Flowering genes: No pathway named

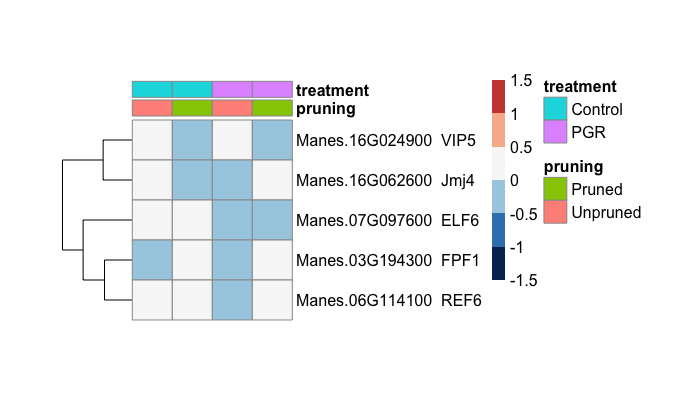

h) Flowering genes: Photoperiod and Circadian rhythm

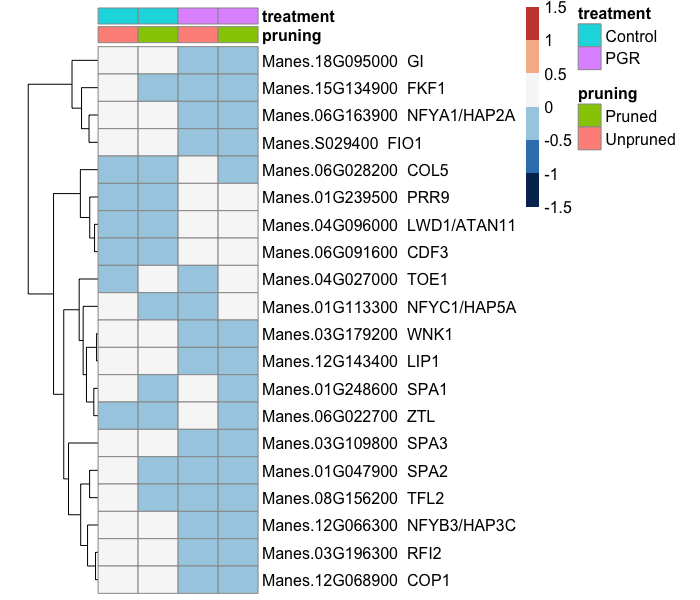

i) Flowering genes: Autonomous

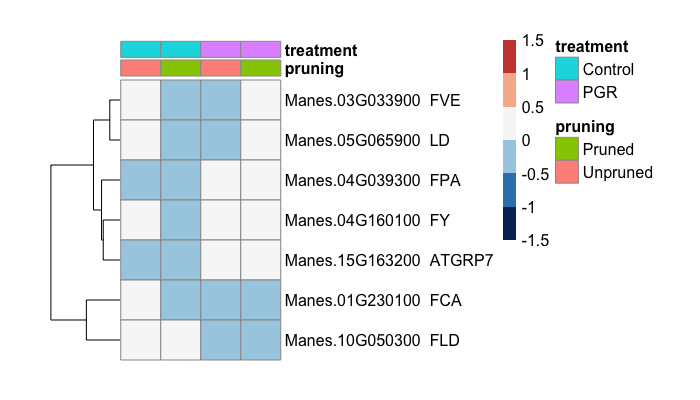

**Supplementary Figure 3** Flowering Genes by pathways

Scale

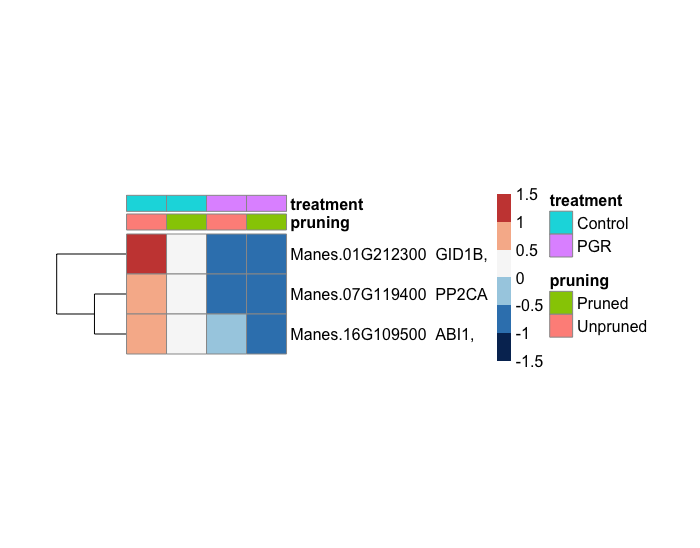

a) Auxin

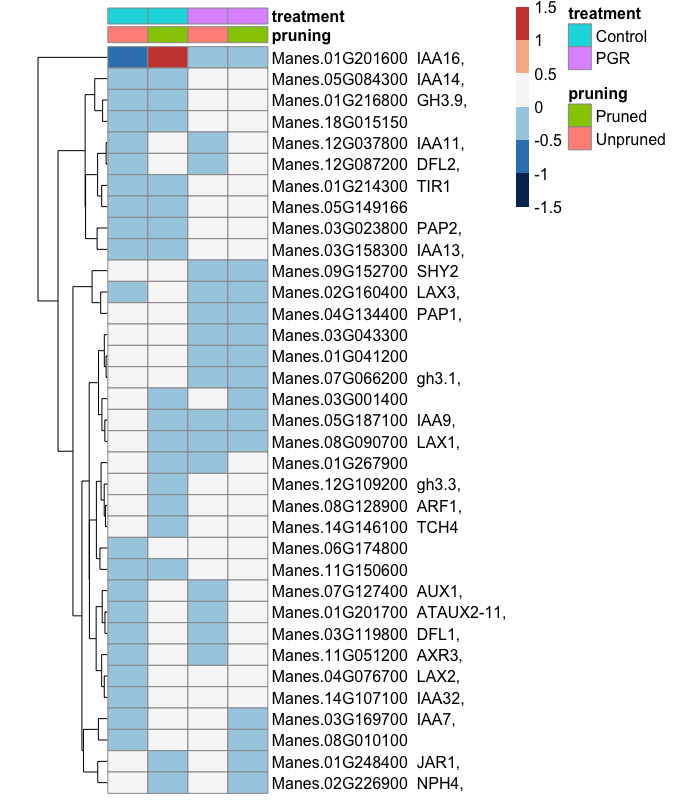

b) ABA

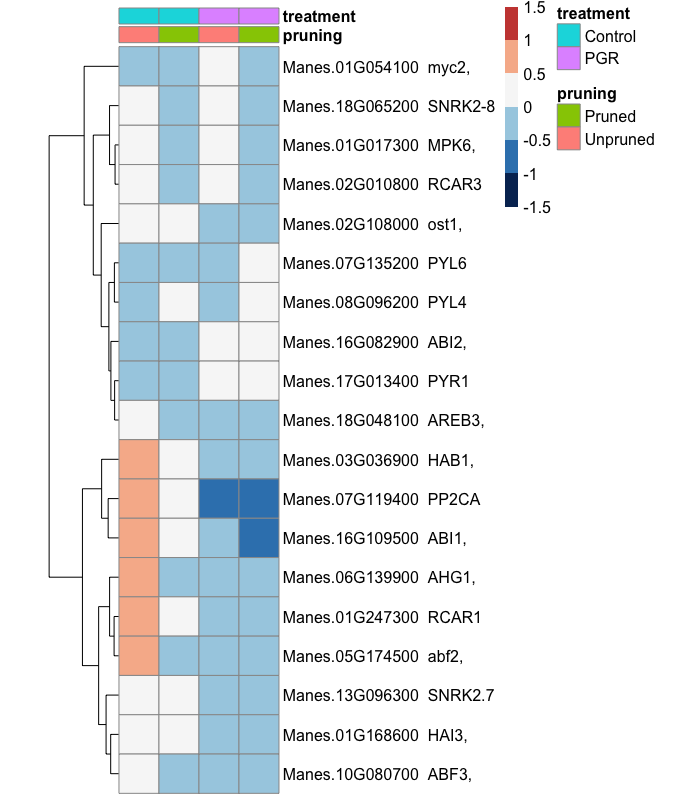

c) GA

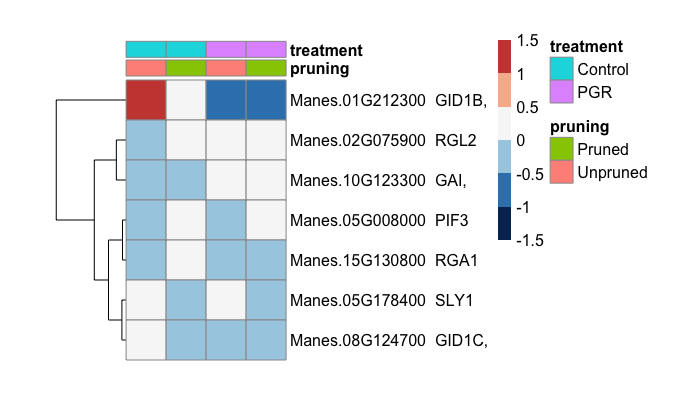

d) Ethylene

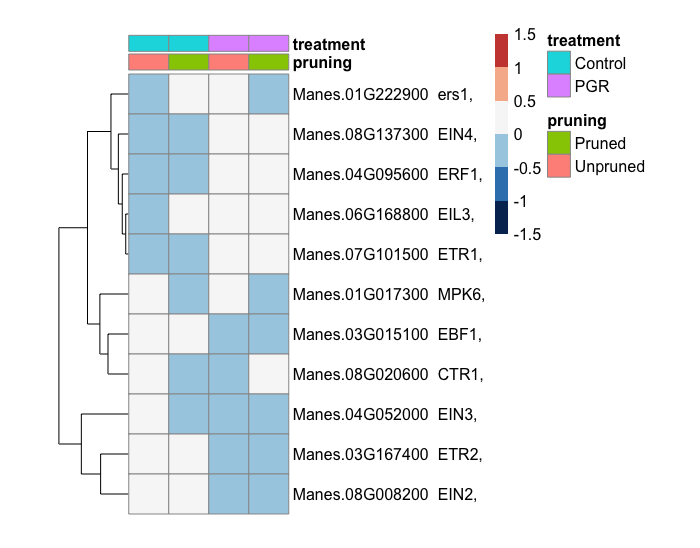

e) Cytokinin

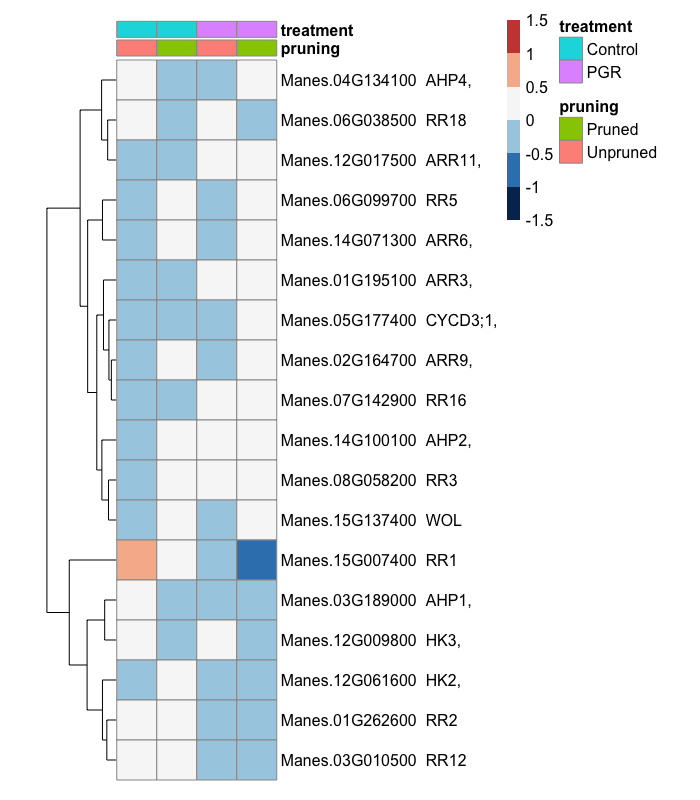

f) BR

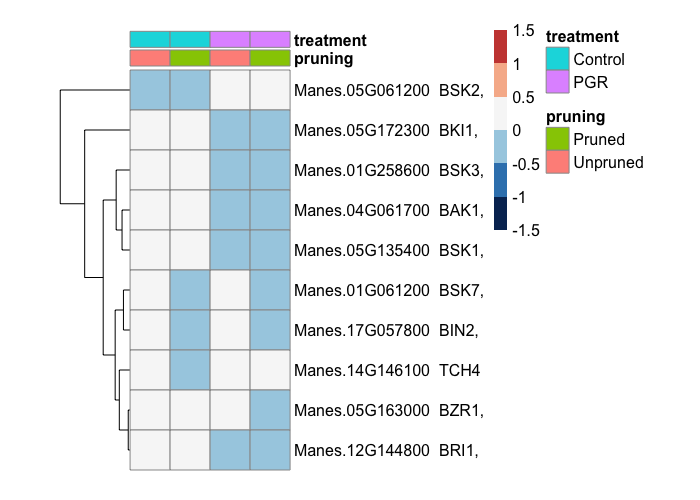

g) JA

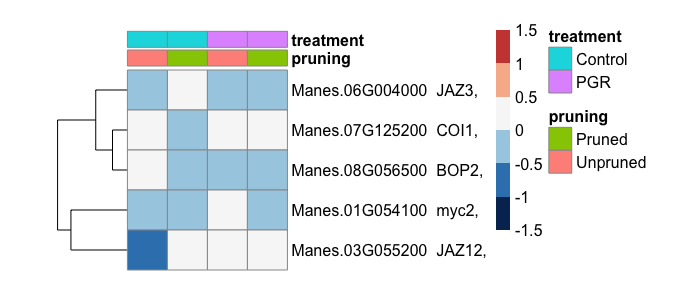

h) SA

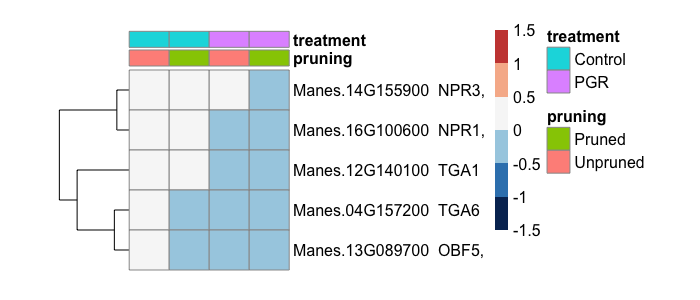

**Supplementary Figure 4** Select hormonal signaling genes by hormonal pathways

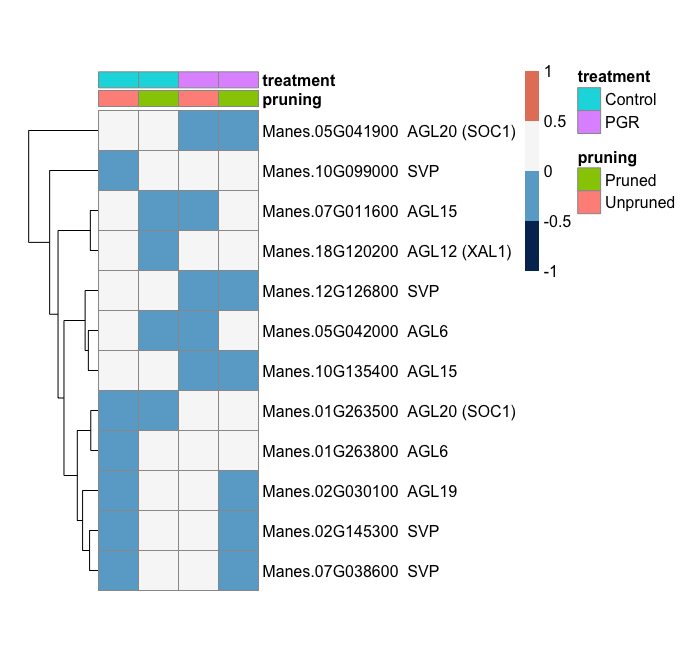

A-Class genes

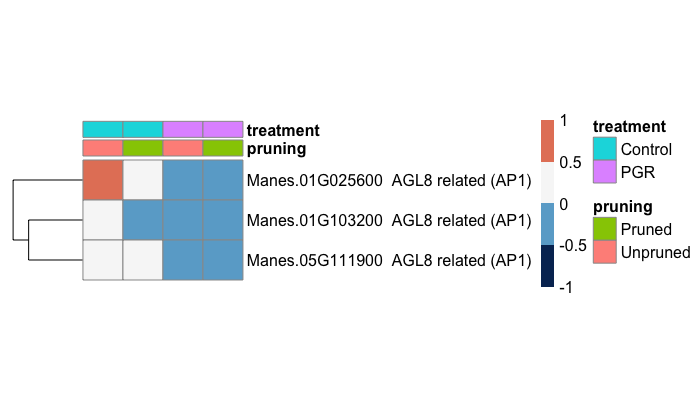

B-class genes

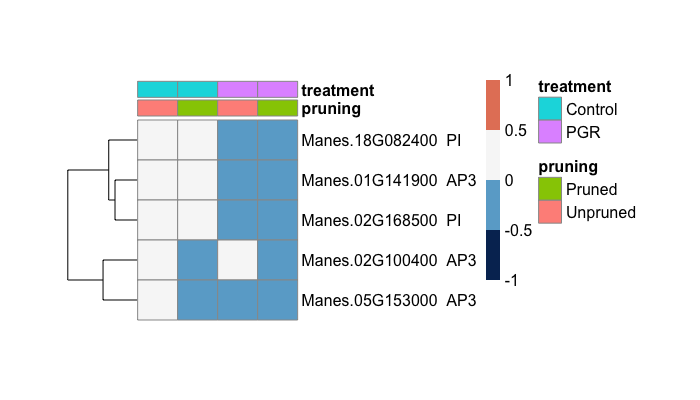

C-class genes

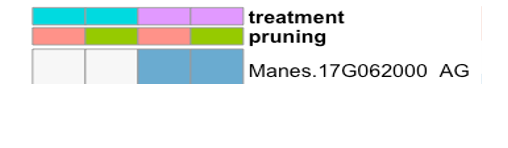

D-Class genes

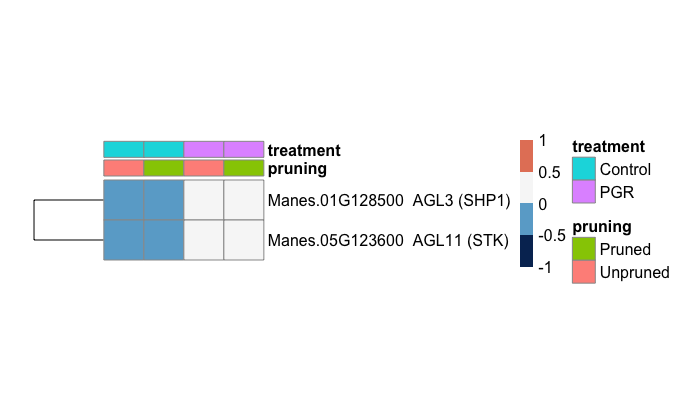

E-Class genes

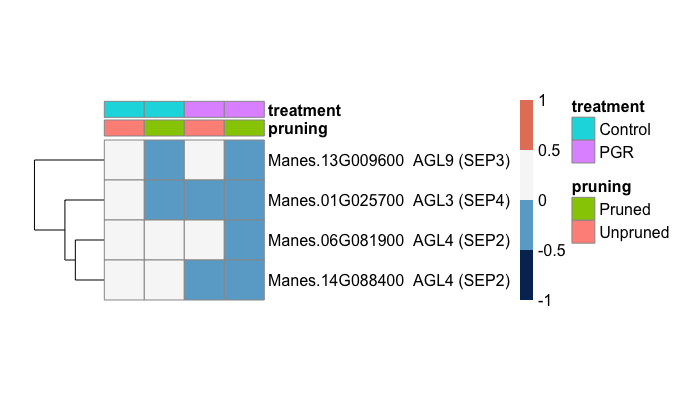

**Supplementary Figure 5** Flowering regulatory genes that were differentially expressed (P≤0.05) in response to PGR and pruning treatments. Shown are genes in each class of MADS-Box transcription factors in the ABCDE model for flower developmental regulation.
